# Probiotic treatment causes sex-specific neuroprotection after traumatic brain injury in mice

**DOI:** 10.1101/2024.04.01.587652

**Authors:** Morgan Holcomb, Austin Marshall, Hannah Flinn, Mariana Lozano, Sirena Soriano, Fernando Gomez-Pinilla, Todd J. Treangen, Sonia Villapol

**Author notes:** Corresponding author: Sonia Villapol, Ph.D. Center for Neuroregeneration and Department of Neurosurgery, Houston Methodist Research Institute, 6670 Bertner Avenue, R11-212, Houston, TX 77030, USA.

## Abstract

**Background:** Recent studies have shed light on the potential role of gut dysbiosis in shaping traumatic brain injury (TBI) outcomes. Changes in the levels and types of *Lactobacillus* bacteria present might impact the immune system disturbances, neuroinflammatory responses, anxiety and depressive-like behaviors, and compromised neuroprotection mechanisms triggered by TBI.

**Objective:** This study aimed to investigate the effects of a daily pan-probiotic (PP) mixture in drinking water containing strains of *Lactobacillus plantarum, L. reuteri, L. helveticus, L. fermentum, L. rhamnosus, L. gasseri,* and *L. casei*, administered for either two or seven weeks before inducing TBI on both male and female mice.

**Methods:** Mice were subjected to controlled cortical impact (CCI) injury. Short-chain fatty acids (SCFAs) analysis was performed for metabolite measurements. The taxonomic profiles of murine fecal samples were evaluated using 16S rRNA V1-V3 sequencing analysis. Histological analyses were used to assess neuroinflammation and gut changes post-TBI, while behavioral tests were conducted to evaluate sensorimotor and cognitive functions.

**Results:** Our findings suggest that PP administration modulates the diversity and composition of the microbiome and increases the levels of SCFAs in a sex-dependent manner. We also observed a reduction of lesion volume, cell death, and microglial and macrophage activation after PP treatment following TBI in male mice. Furthermore, PP-treated mice show motor function improvements and decreases in anxiety and depressive-like behaviors.

**Conclusion:** Our findings suggest that PP administration can mitigate neuroinflammation and ameliorate motor and anxiety and depressive-like behavior deficits following TBI. These results underscore the potential of probiotic interventions as a viable therapeutic strategy to address TBI-induced impairments, emphasizing the need for gender-specific treatment approaches.

## BACKGROUND

Traumatic brain injury (TBI) remains a significant cause of death and disability worldwide [1]. The long-term effects of TBI are associated with an increased risk of neurodegenerative and neuropsychiatric disorders, such as Alzheimer’s and Parkinson’s disease [2, 3]. TBI leads to severe outcomes, with no existing treatments for neuroprotection or regeneration. The pathophysiology of TBI is closely linked to neuroinflammation, predominantly driven by microglial cells [4], and infiltrated macrophages, neutrophils, and leukocytes that contribute to secondary brain damage post-TBI [5]. The inflammatory responses within the central nervous system (CNS) can lead to alterations in peripheral organs. For instance, inflammation in the intestinal lining can trigger an imbalance in the gut microbiome, known as gut dysbiosis [6].

Recent research suggests that many symbiotic bacteria living in our intestines may substantially influence how the brain responds to chronic inflammation and disease [7]. The microbiome-gut-brain axis represents a bidirectional connection between the gut, microbial communities, and brain. This connection is facilitated through the bloodstream or vagus nerve [8], affecting neurological health [9] and the pathogenesis of neurological disorders [10]. The microbiome plays a critical role in essential neurological processes and structures, such as maintaining the integrity of the blood-brain barrier (BBB) [11], myelin formation [12], and microglial activation [13].

Interestingly, survivors of TBI often report gastrointestinal complaints, and animal studies have demonstrated gastrointestinal dysfunction and dysbiosis following TBI [10]. Our previous studies have established a link between TBI and changes in gut microbiota, particularly a disruption known as bacterial dysbiosis [14]. Imbalances in the gut microbiota are associated with worsened outcomes of TBI by leading to neurodegeneration [15] and negative impacts on brain recovery [16]. Alterations in the gut microbiome extend beyond immediate local impacts, affecting essential bacterial populations such as the beneficial *Lactobacillaceae* family, which are vital for maintaining gut health and are prevalent within the human gastrointestinal system [17]. A decrease or lack of these advantageous microbes can amplify neuroinflammation and exacerbate neurocognitive impairments linked to TBI.

Furthermore, TBI has also been associated with decreased levels of microbial metabolites, such as short-chain fatty acids (SCFAs), negatively influencing the healing process [18]. Recent research in animal models indicates that altering the gut microbiome via fecal microbiota transplants can improve recovery outcomes from TBI [19], and stroke [20]. These benefits are thought to be associated with the production of SCFAs, including butyrate, propionate, and acetate, generated through the bacterial breakdown of dietary fibers [21, 22]. SCFAs are key in diminishing inflammation and mitigating oxidative stress [23], impacting BBB integrity, stimulating neurogenesis, reducing neuroinflammation, and promoting microglial maturation [24, 25]. Furthermore, probiotic treatments, which involve administering live beneficial microorganisms like *Lactobacillus* strains, have proven effective in boosting SCFA levels [26, 27] thus potentially offering significant health advantages to the host [28].

Despite these previous findings, there has been no investigation into the effects of probiotics comprising *Lactobacillus* strains on the modulation of microglia and motor and cognitive functions by altering the gut microbiota composition in a mouse model of TBI. We tested the hypothesis that administrating a mixture of probiotics from the *Lactobacillus* strains following TBI in mice would reduce gut dysbiosis and improve motor function by attenuating the neuroinflammatory response. Our findings support the idea that *Lactobacillus spp.* treatment could represent a promising therapeutic approach to managing TBI, which would be easy to translate into a clinical setting, considering probiotics’ well-documented safety and tolerability.

## MATERIALS AND METHODS

### Animals, traumatic brain injury model, and probiotic administration

For our experiments, we utilized male and female C57BL/6 mice, aged 9-12 weeks and weighing 20-26 g, obtained from (Jackson Laboratories, Bar Harbor, ME, US). The mice were housed in an environment with a 14-h light/10-h dark cycle at a stable temperature of 23 °C ± 2 °C and had continuous access to food and water. After arriving, the mice were given a minimum of three days to adjust to their new surroundings before participating in any experimental procedures. Both adult male and randomly cycling female C57BL/6 mice were divided into groups for the study. We included only males for the germ-free (GF) mice because males have a higher inflammatory response after TBI than females [29]. C57BL/6-GF male mice (12 weeks) were obtained from the Baylor College of Medicine Gnotobiotic Rodent Facility (Houston, TX). An internal standard was serially diluted to assess the “*germ-free*” status of the mice upon arrival, and the copy number of the 16S rRNA gene in feces from each transfer crate was analyzed using quantitative PCR (qPCR). 16S rRNA gene was not detected in feces from transfer crates, confirming that the mice were GF.

Anesthesia was induced with 4-5% isoflurane, then maintained at 1.5-2% with a 1-1.5 L/min oxygen flow rate. Each mouse, under anesthesia, was secured in a stereotaxic frame for the surgery. A moderate to severe TBI was inflicted on the left side of the brain, targeting the primary motor and somatosensory cortices, using an electromagnetic device (Leica StereoOne Impactor) driven controlled cortical impact (CCI) injury. Throughout the surgical procedure, the body temperature of the mouse was maintained at 37 °C using a heating pad. A midline incision approximately 10 mm long was made on the skull. The skin and fascia were pulled back to expose the skull, where a 4 mm craniotomy was then performed on the left parietal bone. The sterilized tip of 1.5 mm diameter was placed onto the surface of the exposed dura and set to deliver an impact to the cortical surface with a velocity of 3.25 m/s and a tissue deformation of 1.5 mm. Mice in the sham-operated group underwent identical procedures, including isoflurane anesthesia, but without the CCI injury. Post-TBI, the incision was sutured, and the mouse was placed in a warmed recovery chamber to maintain average body temperature. All animals were closely monitored for 4 h post-surgery and then checked daily. The mice were distributed into four distinct groups, each consisting of 4 to10 mice: (i) a sham group (sham-operated control, vehicle (VH)-treated), (ii) a sham group (sham-operated with pan-probiotic (PP) treatment), (iii) a CCI group (VH-treated), and (iv) a CCI group (PP treatment) (**Fig. 1**).

**Figure 1.**
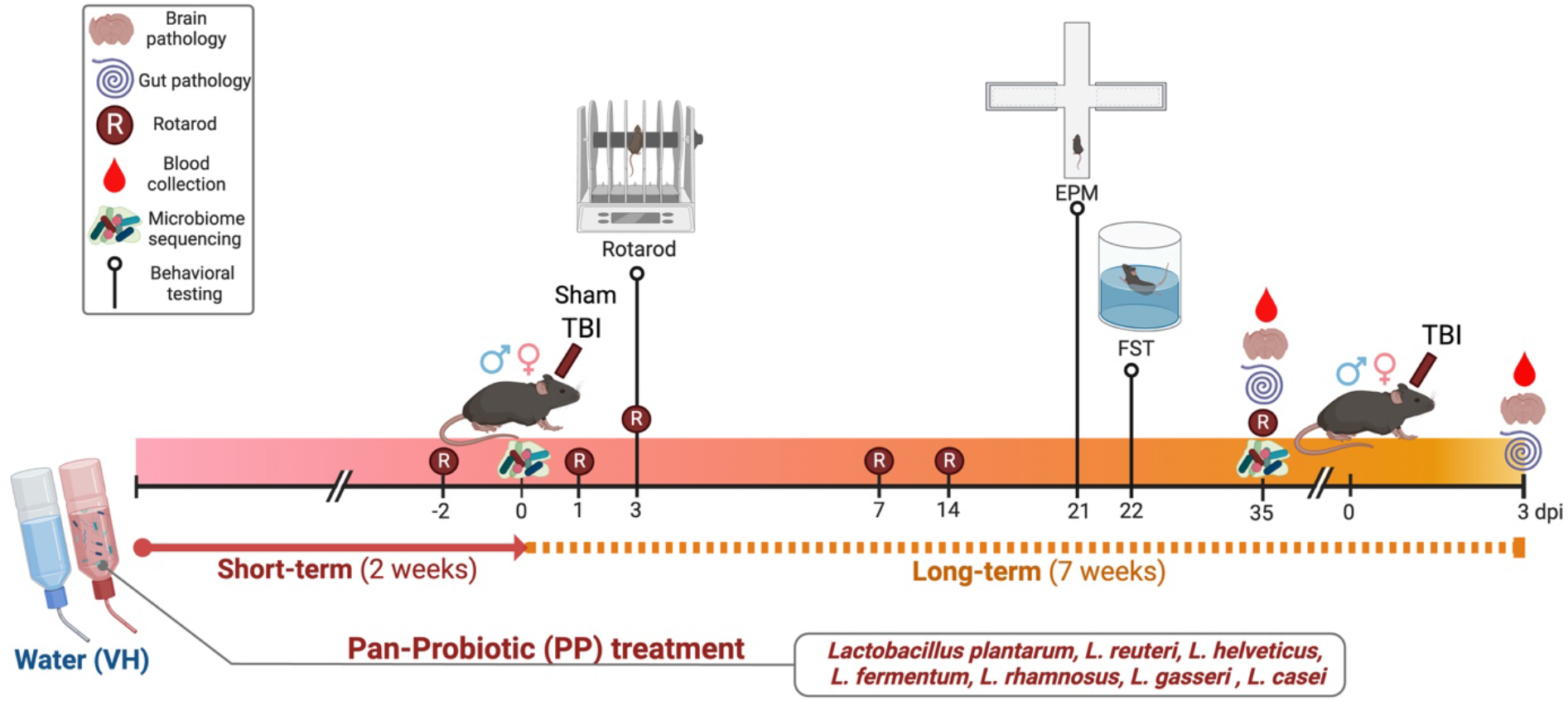
Timeline and Evaluation of Probiotic treatment in a Mouse Model of Traumatic Brain Injury (TBI). Mice received pan-probiotic treatment for 2 weeks before traumatic brain injury (TBI) and continued treatment for up to 7 weeks. Motor skills were assessed at various intervals using the Rotarod test (R), and blood samples were collected to examine metabolites. Additionally, fecal samples were obtained for microbiome sequencing. Behavioral assays, including the Elevated Plus Maze (EPM) and Forced Swim Test (FST), were employed to gauge anxiety and depressive-like symptoms. Immunohistochemical examination of brain tissues was conducted at acute (3 days post-TBI) and chronic (35 days post-TBI) treatments.

Pan-probiotic (PP) mixture in drinking water containing strains of *Lactobacillus plantarum, L. reuteri, L. helveticus, L. fermentum, L. rhamnosus, L. gasseri,* and *L. casei*, administered for either two or seven weeks before inducing TBI on both male and female mice. Mice were treated with PP in their drinking water for 2 weeks before TBI and continued for 5 more weeks until euthanasia for tissue collection at 35 days post-injury (dpi). A separate set of mice was treated with PP for 7 weeks, injured and then euthanized at 3 dpi to assess the acute effects. All mice in the four groups were given a standard diet and either PP-treated or regular drinking water for VH groups. For euthanasia, the mice were deeply anesthetized using isoflurane at either 3 dpi (acute injury) or 35 dpi (chronic injury) and underwent transcardiac perfusion with cold phosphate-buffered saline (PBS) followed by 4% paraformaldehyde. The GF mice were removed from the isolator in the gnotobiotic facility and transferred to the laminar flow with high efficiency particulate air (HEPA) filter to perform the CCI injury model. After surgery, GF mice were transferred to autoclaved cages, provided with autoclave chow and water, and maintained in a conventional animal holding room for 3 days until euthanasia. Brain tissues were then harvested for immunohistochemical analysis, and serum samples were collected via centrifugation of blood and stored at −80 °C for SCFAs analysis. Samples of intestinal tissue were promptly collected and immediately frozen in dry ice. This study was performed under the National Institute of Health guidelines, and the Institutional Animal Care approved all experiments and Use Committees at Baylor College of Medicine and Houston Methodist Research Institute (Houston, TX).

### Preparation of bacterial cell suspension for oral feeding

The PP mixture comprised various *Lactobacillus* strains supplied by the American Type Culture Collection (ATCC) (Manassas, VA, USA). The strains included *Lactobacillus gasseri* (ATCC 33323), *Lactobacillus plantarum* (ATCC BAA-793), *Lactobacillus reuteri* (ATCC 23272), *Lactobacillus helveticus* (ATCC BAA-2840), *Lactobacillus fermentum* (ATCC 23271), *Lactobacillus rhamnosus* (ATCC BAA-2836), and *Lactobacillus casei* (ATCC BAA-2843). Following the supplier’s protocol, we cultured the strains in MRS broth (Beckton Dickinson, Sparks, MD), and preserved them as glycerol stocks at -80°C. To initiate culture growth, we prepared 5 mL of pre-warmed MRS broth with the glycerol stocks and then incubated these starter cultures at 37°C in a 5% CO2 environment for 2 h. The bacteria were cultured overnight in 1 L of MRS broth under identical conditions until they reached the logarithmic growth phase, confirmed by measuring optical density at 600 nm (OD600). After growth, the bacterial cells were collected through several centrifugation steps at 3000xg and 4°C, followed by a wash in cold PBS. We then resuspended the bacterial pellets in 10% glycerol in PBS and promptly froze them in 1 mL aliquots for future administration to mice. The viability and concentration of the bacteria were verified by serial dilution and colony-forming unit (CFU) enumeration on agar plates.

### Brain Tissue Preparation and Quantification of Lesion Volume

The brains were extracted and further fixed in 4% paraformaldehyde overnight. After 24 h of fixation, the brains underwent cryopreservation in a 30% sucrose solution. Coronal brain sections measuring 16 μm in thickness were cut using a cryostat (Epredia Cryostar NX50, Fisher Scientific, Waltham, MA). The sectioning process was done in intervals of 0.5 mm throughout the brain, beginning 1.56 mm anterior from the bregma landmark to ensure a consistent and thorough examination of the affected areas for subsequent histochemical analysis. An average of 10-12 brain sections, equally spaced from 0 to -2.70 mm relative to the bregma, were selected for cresyl violet staining to target the area affected by injury. Sections were placed on gelatin-coated glass slides (SuperFrost Plus, Thermo Fisher Scientific, IL) and submerged in 0.5% cresyl violet solution (Sigma-Aldrich, St. Louis, MO), prepared in distilled water and filtered. Following staining, the slides underwent a dehydration process in graded ethanol solutions (100%, 95%, 70%, and 50%) for 2 min each and were then cleared in xylene twice, each for 2 min. The sections were then covered and slipped using a Permount mounting medium (Thermo Fisher Scientific) for preservation. The lesion area was quantified by examining every 16^th^ section throughout the full range of the lesion, using a methodology outlined in our previous publications [29–32]. The area of the ipsilateral hemisphere was also measured for each corresponding section. The lesion volume was calculated by multiplying the aggregate of lesion areas by the interval between sections. To determine the percent lesion volume, we divided the lesion area by the area of the total ipsilateral hemisphere.

### Serum SCFA analysis

SCFAs were analyzed by derivatization procedure. 40 µL of serum was added to 40 µL of acetonitrile, vortexed, and centrifuged. 40 µL of the supernatant, 20 µL of 200 mM 12C6- 3- Nitrophenylhydrazine (3NPH), and 120 mM 1-Ethyl-3-(3-dimethylaminopropyl)carbodiimide (EDC) were combined. 20 µL of hydrochloric acid was added and incubated for 30 min at 40°C. The resulting mixture was cooled and made up to 1.91 mL with 10% aqueous acetonitrile. 5 µL of the sample was injected into LC/MS/MS. SCFAs were separated using mobile phases 0.1% formic acid in water (mobile phase A) and 0.1% formic acid in acetonitrile (mobile phase B). Separation of metabolites was performed on Acquity UPLC HSS T3 1.8 um (2.1×100mM). The SCFA were measured in ESI negative mode using a 6495 triple quadrupole mass spectrometer (Agilent Technologies, Santa Clara, CA) coupled to an HPLC system (Agilent Technologies, Santa Clara, CA) with multiple reaction monitoring (MRM). The acquired data was analyzed using Agilent Mass Hunter quantitative software. (Agilent Technologies, Santa Clara, CA).

### Immunofluorescence analysis and cell death assay

We prepared serial free-floating brain sections, each 16 μm thick, for our immunohistochemical studies, at the dorsal hippocampus level. These sections underwent immunohistochemistry processing, which included 3 successive 5-min washes in PBS enhanced with 0.5% Triton X-100 (PBS-T). Sections were then treated with 5% normal goat serum (NGS) in PBS-T to block nonspecific binding for 1 h at room temperature. Overnight incubation at 4°C followed, using 3% NGS in PBS-T with primary antibodies targeting anti-rabbit Iba-1 (Wako, #019-19741) at a dilution of 1:500 for labeling microglia/macrophages, and anti-rat F4/80 (R&D Systems, #MAB5580) at a dilution of 1:200. The following day the sections were washed 3 times for 5 min each in PBS-T and incubated with the corresponding secondary antibodies (all 1:1000, Invitrogen), for 2 h at room temperature. The sections were then rinsed with PBS three times for 5 min each, and incubated in PBS with DAPI solution (1:50,000, Sigma-Aldrich, St. Louis, MO) for counterstained nuclei. The sections were rinsed with distilled water and covered with Fluoro-Gel with Tris Buffer mounting medium (Electron Microscopy Sciences, Hatfield, PA). Semiquantitative image analysis of the staining’s in the cortical regions was performed using Image J software as previously described [29]. To assess cell death, brain sections were processed for DNA strand breaks using Terminal deoxynucleotidyl transferase dUTP nick end labeling (TUNEL) from Fluorescence *In Situ* Cell Death Detection kit (Roche Diagnostic, Indianapolis, IN) according to the manufacturer’s instructions.

### Image Acquisition and Sholl Analysis of Microglia Morphology

To analyze the microglia morphology in the injured brains after 3 dpi, we used confocal microscopy to collect full thickness 40X magnification z-stack images using a Leica confocal microscope at the injured cortex from 16 um-thick brain sections that were immunostained for Iba-1 (n=4 per group). Z-stacks were collected at 2048 X 2048 resolution with 3 frame averages for each color channel and were exported as .TIF files. After importing the Z-stacks, we performed Sholl analysis of microglial morphology using the Neurolucida Explorer (MBF Bioscience) setting increments of 2 µm with a starting radius of 2 µm, and the branches were detected using the user-guided tree tracing. When examining microglial processes in detail using Sholl analysis, the number of intersections and the average length of processes were quantified across different distances from the cell body. We combined intersection data from each microglia in a region into single profile plots, enabling comparison of the overall morphology between cortical microglia in the injured brains treated with VH or PP.

### Histological Analysis of the Gut

To assess gut inflammation, we evaluated intestinal villus-to-crypt ratios in the jejunal region of the intestine. The intestines harvested from mice treated with VH or PP after 3 or 35 dpi were initially fixed in 4% paraformaldehyde for 48 h and transferred to 70% ethanol. Tissue processing was carried out using a Shandon Excelsior ES Tissue Processor, and embedding took place on a Shandon HistoCentre Embedding System, following the manufacturer’s standard protocols for processing and embedding. Subsequently, slides were sectioned to a thickness of 5 μm. The paraffin sections were then subjected to deparaffinization and rehydration. Hematoxylin solution was applied for 6 h at a temperature ranging from 60 to 70°C, followed by rinsing with tap water until the water became colorless. The tissue was differentiated using 0.3% acid alcohol in water, with 2 min treatments. After rinsing with tap water, the sections were stained with eosin for 2 min. Alcian blue staining was employed to visualize mucins produced by the goblet cells in the intestine. Deparaffinized and rehydrated sections were immersed in Alcian blue solution for 30 minutes and then counterstained with 0.1% Nuclear Fast Red for 5 minutes. Finally, the intestine sections were dehydrated and mounted using permanent mounting media.

### Quantitative Analysis

For quantitative analysis of immunolabeled sections, we implemented unbiased, standardized sampling techniques to measure tissue areas corresponding to the injured cortex showing positive immunoreactivity, as we previously described [29, 31]. For proportional area measurements, the microglia/macrophages Iba-1-immunoreactivity was reported as the proportional area of tissue occupied by immunohistochemical stained cellular profiles within a defined target area. Thresholded images converted to 8-bit grayscale were made using ImageJ (NIH, Bethesda, MD). The thresholding function was then used to set a black-and-white threshold corresponding to the imaged field, with the averaged background subtracted. Once a threshold was set, the “*Analyze Particles*” function was used to sum up the total area of positive staining and calculate the fraction of the total positive area for the stain as previously described [29]. To quantify the number of TUNEL, Iba-1, and F4/80+ cells in the injured cortex, an average of four coronal sections from the lesion epicenter (−1.34 to −2.30 mm from Bregma) were counted and imaged for each animal, *n* = 8/group.

### Collection and Extraction of Fecal Samples for DNA Analysis

Fresh stool pellets were aseptically collected and placed in sterile tubes, immediately snap-frozen, and subsequently stored at −80°C for preservation. Genomic bacterial DNA was extracted from these frozen stool samples utilizing the QIAamp PowerFecal Pro DNA Kit (Qiagen, Germantown, MD). To facilitate DNA extraction, bead beating was carried out in three cycles, each lasting one min, at a speed of 6.5 m/s. There was a 5 min rest period between each cycle. This mechanical disruption was performed using a FastPrep-24 system (MP Biomedicals, Irvine, CA). The DNA isolation proceeded per the manufacturer’s instructions, following the bead-beating process. The concentration of the extracted genomic DNA was subsequently quantified using a DS-11 Series Spectrophotometer/Fluorometer (DeNovix, Wilmington, DE).

### Sequencing of 16S rRNA V1-V3 regions

The specific gene sequences employed were designed to target the 16S ribosomal RNA gene region V1 to V3. The primers used for amplification contain adapters for MiSeq sequencing and single-index barcodes so that the PCR products may be pooled and sequenced directly, targeting at least 10,000 reads per sample [33]. Primers used for the 16S V1-V3 amplification were 27F (AGAGTTTGATYMTGGCTCAG, where Y = C (90%) or T (10%); M = A (30%) or C (70%)) and 534R (ATTACCGCGGCKGCTGG, where K = G (10%) or T (90%)) [34]. Amplicons were generated using primers corresponding to the variable regions, and the PCR products were purified. Subsequently, sequencing libraries for the V1-V3 target were constructed following the instructions provided by the Illumina MiSeq system with end products of 300 bp paired-end libraries.

### Amplicon sequence analysis pipeline

Raw data files in binary base call (BCL) format were converted into FASTQs and demultiplexed based on the single-index barcodes using the Illumina ‘bcl2fastq’ software. Demultiplexed read pairs underwent an initial quality filtering using bbduk.sh (BBMap version 38.82), removing Illumina adapters, PhiX reads and reads with a Phred quality score below 15 and length below 100 bp after trimming. 16S V1-V3 quality-controlled reads were then merged using bbmerge.sh (BBMap version 38.82), with merge parameters optimized for the 16S V1-V3 amplicon type (vstrict=t qtrim=t trimq=15). Further processing was performed using nf-core/ampliseq version 2.8.0 of the nf-core collection of workflows, utilizing reproducible software environments from the Bioconda and Biocontainers projects [35–38]. Data quality was evaluated with FastQC (version 0.12.1) and summarized with MultiQC (version 1.18) [39]. Sequences were processed sample-wise (independent) with DADA2 (version 1.28) to eliminate PhiX contamination, trim reads (forward reads at 275 bp and reverse reads at 265 bp; reads shorter than this were discarded), discard reads with > 2 expected errors, to correct errors, to merge read pairs, and to remove PCR chimeras. Ultimately, 3880 amplicon sequencing variants (ASVs) were obtained across all samples [40]. Between 29.81% and 44.06% of reads per sample (average 36.8%) were retained. The ASV count table contained 2337653 counts, at least 9201 and 34821 per sample (average 19980). Taxonomic classification was performed by DADA2 and the database ‘Silva 138.1 prokaryotic SSU’ [41]. ASV sequences, abundance, and DADA2 taxonomic assignments were loaded into QIIME2 (version 2023.7) [42]. Within QIIME2, the final microbial community data were collected into an .rds file found within the phyloseq (version 1.44) folder of the nf-core ampliseq output [43]. The resulting phyloseq data frame object was merged with the phylogenetic tree calculated within QIIME2. The phyloseq object was used in the creation of alpha and beta diversity plots, PERMANOVA calculations with vegan::adonis2 (version 2.6-5), relative abundance bar plots with microViz (version 0.12.1), nextflow, and R (version 4.3.2) scripts, and differentially abundant taxa calculations using ANCOMBC2 [44].

### Western Blot Analysis

Serum was collected by allowing trunk blood to clot at room temperature for 30 min, followed by centrifugation at 4000 revolutions per minute (rpm) and 4°C for 20 min. The serum was then stored at −80 °C. For analysis, serum samples were diluted 1:5 with Laemmli sample buffer (Bio-Rad Laboratories, Hercules, CA) and heated at 100 °C for 10 min. The samples were then loaded onto 12% Mini-PROTEAN TGX Stain-Free gels (Bio-Rad Laboratories, Hercules, CA) for protein separation. Following electrophoresis, proteins were transferred to nitrocellulose membranes (Bio-Rad Laboratories, Hercules, CA). The membranes underwent blocking with 5% w/v skim milk powder in PBS-Tween 20 (PBS-Tw) for 1 h at room temperature. They were then incubated overnight at 4°C with a goat anti-SAA primary antibody (1:500; AF2948, R&D Systems, Minneapolis, MN). This was followed by a 1 h room temperature incubation with a horseradish peroxidase-conjugated rabbit anti-goat secondary antibody (1:3000; Thermo Fisher Scientific, Waltham, MA). The membranes were developed using Clarity Western ECL (Bio-Rad Laboratories, Hercules, CA) and imaged with a ChemiDoc MP system (Bio-Rad Laboratories, Hercules, CA). Densitometry quantification and chemiluminescence imaging were carried out using ImageLab 6.0.1 software (Bio-Rad Laboratories, Hercules, CA).

### Behavioral tests. Rotarod Test

The rotarod test is a method employed to assess sensorimotor function in mice [45]. For this evaluation, we used the Rotamex 5 system from Columbus Instruments (Columbus, OH) and followed the protocol as we have previously described [30, 46]. Each mouse underwent a training regimen spanning 2 days, with 3 daily trials. The procedure began by initially placing the mouse on a stationary rod, allowing the animal to explore for 30 sec. Then, the rod began to rotate, with the drum’s speed gradually increasing from 4 to 40 rpm. Each trial concluded when the mouse fell off the rotarod, or a 5-min time span had elapsed, and the latency to fall was recorded. The mean value of these measurements served as the baseline assessment for each mouse. The testing sessions were conducted at 1, 3, 7, 14, and 35 dpi, with two investigators blinded to the animal groups overseeing the assessments.

### Elevated Plus Maze Test

The Elevated Plus Maze Test (EPM) is a neurobehavioral paradigm that examines the conflict between an mouse’s innate exploratory drive and its fear of open areas. The EPM is designed to measure anxiety-related behaviors. It consists of a plus-shaped apparatus with 2 open and 2 closed arms. Mice typically avoid open spaces due to the increased risk of exposure to predators. Therefore, animals that spend more time in the open arms are considered to exhibit less anxiety-like behavior. In our study, we utilized an EPM apparatus based on the design introduced by Pellow et al. [47]. The plus-shaped maze is custom-made using white acrylic material. It comprises 2 open arms (50×10 cm) and 2 closed arms (50×10×40 cm), positioned in such a way that the 2 arms of each type are situated opposite to each other and connected by a central platform (5×5 cm). The maze is elevated to a height of 75 cm above the floor. At 21 dpi, each mouse was gently placed at the central platform of the maze, with its head oriented toward one of the open arms. Subsequently, we recorded data pertaining to the time spent in the open arms, over a 5-min duration. These measurements were expressed as percentages. After each trial, the maze’s platform was cleaned with 70% isopropanol. The behavioral assessments of the mice were conducted during the light phase of the light-dark cycle.

### Forced Swimming Test

The Forced Swimming Test (FST) is a widely recognized animal model utilized to study despair behavior and is highly sensitive for evaluating the effects of pharmacological interventions on depression-like behaviors [48]. The FST measures the propensity of the animal to give up when placed in a cylinder filled with water from which they cannot escape. Rodents will initially try to escape, but over time, they will start to float more and struggle less, which is interpreted as a state of behavioral despair akin to depressive-like behavior. In this context, more time spent immobile (floating without trying to escape) is considered an indicator of greater depressive-like behavior. At 22 dpi, mice were placed inside glass cylinders measuring 30 cm in height and 20 cm in diameter, filled with water at a temperature of 25 ± 1°C, reaching a depth of 15 cm. The test duration was established to be 6 min. During the final 4 min of the test, a trained observer, who was blinded to the treatment groups, recorded the behavioral responses of the mice. In this context, despair behavior in animals was quantified in terms of total swimming and immobility. Mouse immobility was passive floating with minimal movement, just enough to keep the animal’s head above the water. The cumulative time spent by the mice in this immobile state served as an indicator of behavioral despair. Additionally, any time the mice spent climbing or swimming was considered part of the total swimming time.

### Statistical Analysis

For the SCFA analysis, differentially expressed metabolites were detected using the unequal variance two-sided Student’s *t*-test followed by Benjamini-Hochberg correction, with significance achieved at an FDR-adjusted *p* < 0.25. A one-way analysis of variance (ANOVA) with post-hoc Tukey’s multiple comparison test was employed for immunohistochemistry data. We adhered to a significance threshold (alpha) 0.05 across all datasets. For the sholl analysis the area under the curve was calculated, and an unpaired Student’s t-test was utilized to analyze mean fluorescence intensity and cell count. We used two-way ANOVA for behavioral analysis followed by post-hoc Tukey’s multiple comparison. Analysis of the rotarod test and histochemical/immunofluorescence utilized a one-way ANOVA post-hoc Tukey’s multiple comparison to compare the time after injury and sex as the independent variables. All mice were randomized to experimental conditions, and experimenters were blind to the treatment groups throughout the study. Data was represented as the mean and standard error of the mean (±SEM), and *p < 0.05, **p < 0.01, ***p < 0.001, and ****p < 0.0001 were considered statistically significant. GraphPad Prism 8 Software (GraphPad; San Diego, CA, US) was used for statistical analysis.

## RESULTS

### Pan-probiotics increase the relative abundance of SCFAs in serum

At 3 dpi, there were no significant differences in the levels of acetate, propionate, butyrate, isobutyrate, 2-methyl-butyrate, isovalerate, valerate, and caproate between VH and PP treatment groups across both male and female mice (**Fig. 2 a-h**). However, there was a general trend for an increase in the butyrate, isobutyrate, valerate, and caproate groups in the PP-treated males compared to females (**Fig. 2 c, d, g, h**). At 35 dpi, notable differences were observed, isobutyrate, 2-methyl-butyrate, isovalerate, valerate, and caproate levels demonstrated significant increases in the PP-treated males compared to VH, with 2-methyl-butyrate and caproate also showing significant increases in PP-treated males versus females (**p < 0.01, *p < 0.05, respectively) (**Fig. 2 l-p**). This indicates that the PP treatment successfully replenishes certain SCFAs, typically diminished following TBI, after 7 weeks of treatment. However, it is worth noting that there was a reduction in the relative abundance of acetate in the female PP mice when compared to their VH counterparts (**Fig. 2i**). These findings suggest that the PP treatment’s effect on SCFA profiles varies by sex in the setting of TBI, potentially influencing metabolic and inflammatory responses.

**Figure 2.**
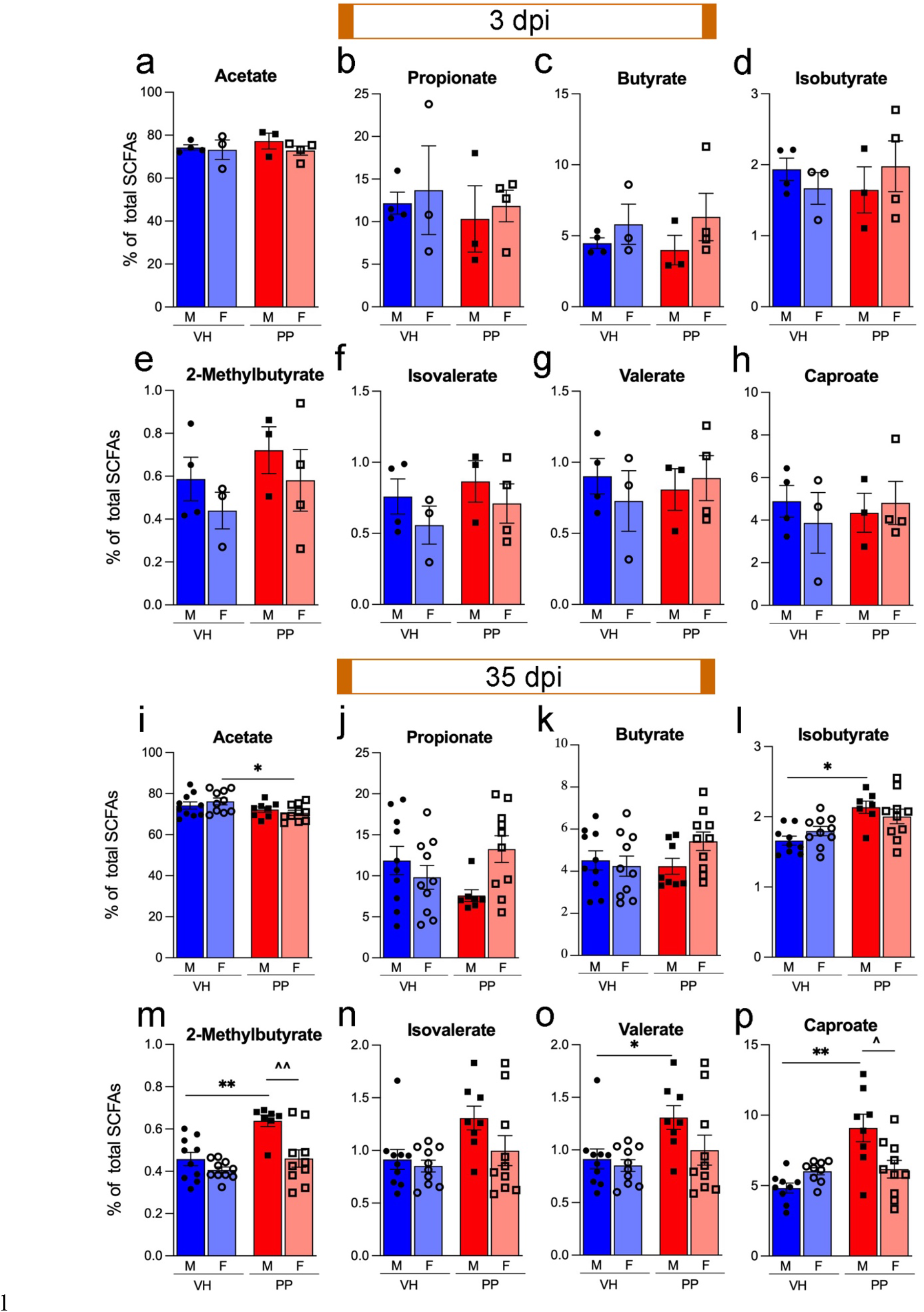
Pan-Probiotic (PP) treatment increases Short-Chain Fatty Acid levels post-injury. (a-h) Initially, at 3 dpi Short-Chain Fatty Acid (SCFA), SCFA levels did not significantly differ. However, there was a notable trend towards higher 2-methyl-Butyrate levels in PP-treated male and female subjects compared to their VH counterparts (e). (i-p) At the 35 dpi, PP-treated male mice displayed significant increases in several SCFAs, including Isobutyrate, 2-methylbutyrate, Valerate, and Caproate (l-p). In contrast, PP-treated female mice experienced a reduction in Acetate levels (i). There were also significant increases at 35 dpi in SCFAs 2-methylbutyrate and Caproate with PP-treated males compared to females (m, p). Sample sizes: 3 dpi VH-treated, PP-treated (n=3-4/group), 35 dpi VH-treated, and PP-treated (n=8-10/group), with statistical significance assessed using a two-sided Student’s *t*-test followed by Benjamini-Hochberg correction, highlighting changes in SCFA metabolites after TBI and probiotic intervention (*p<0.05, **p<0.01, ***p<0.001).

### The effect of pan-probiotic administration on gut microbiota composition

Next, we evaluated whether PP treatment could affect the fecal microbiota composition and abundance based on the number of OTUs in the microbiome. The alpha diversity of the microbiome compositions revealed statistically significant differences between the VH and PP groups in the sham group over a simulated 7-week period, as indicated by the Shannon index (**Fig. 3a**). There was a statistically significant distinction between the VH and PP treatment groups in the TBI mice, according to the Simpson index (**Fig. 3 b**), which adjusts for biases in sample uniformity, leading to a different result compared to the Shannon diversity of the same dataset. The introduction of PP should increase the microbial counts in the gut [49]. The beta diversity analysis of the same study groups shows that there is a statistically significant variation between the VH and PP treatments after 2 weeks (Pr(>F) = 0.014), (R^2^=0.063), (**Fig. 3 e-g**). The relationship between the gut microbiomes of the VH and PP mice was not highly altered at the 2-week timepoint when observing the phylum level, as revealed by the stacked bar representation for the different groups, with a statistical significance between the sham-VH group mice. The Firmicutes to Bacteroidetes ratio showed a significant reduction in the VH group throughout treatment from 2 weeks to the 7 weeks sham-VH group (**Fig. 3 h**), while a significant change was also observed between the sham and TBI groups in females (*p<0.05, **p<0.01, ***p<0.001) (**Fig. 3 i**). No significant differences were detected between the sham and the TBI PP-treated groups, indicating a return towards normal Firmicutes and Bacteroidota levels.

**Figure 3.**
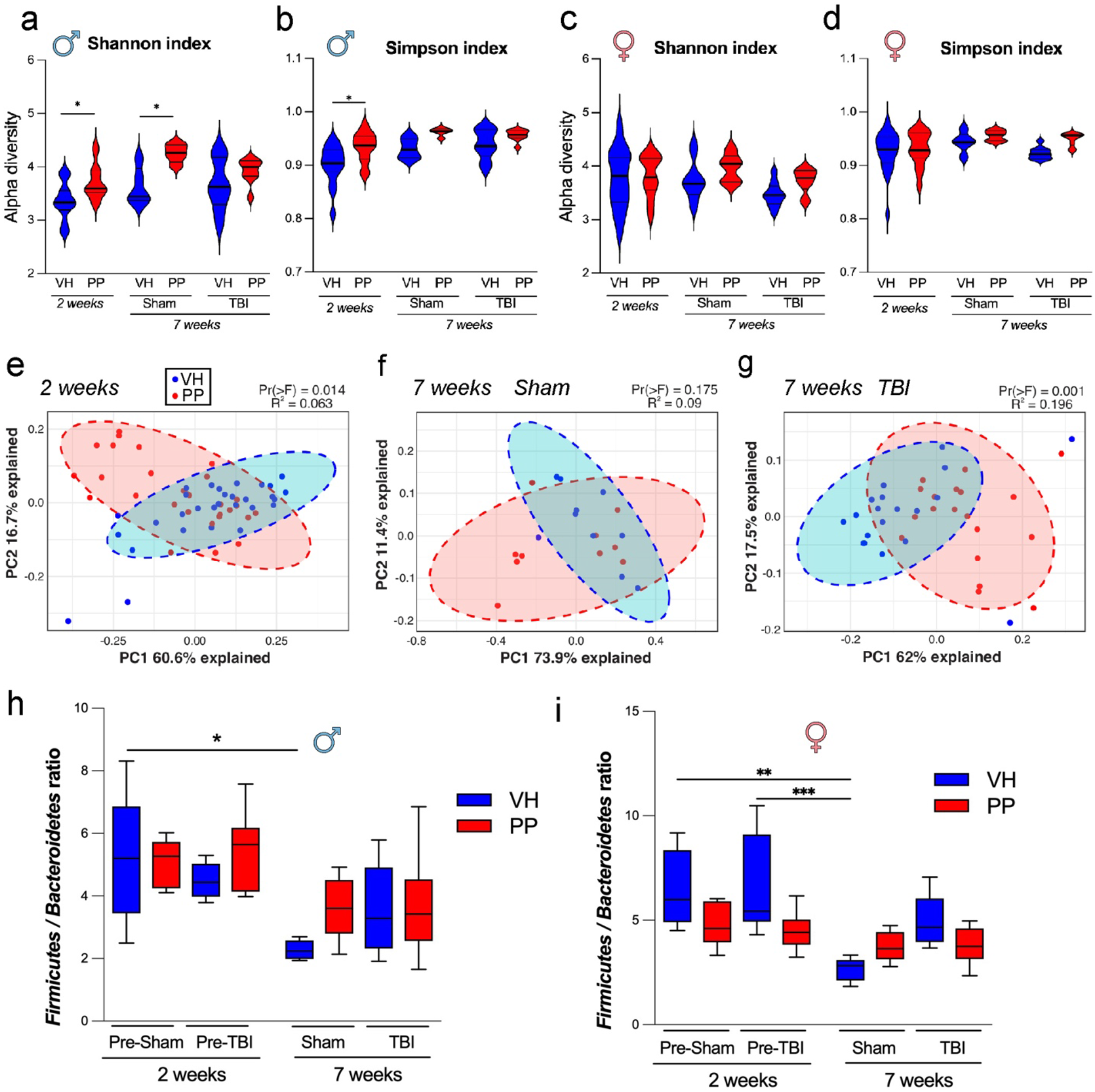
Taxonomic Profile dynamics in the gut microbiome following pan-probiotic treatment. (a-d) The Shannon and Simpson indices in males and females indicate that fecal microbiota diversity was higher following a 2-week pan-probiotic (PP, in red) treatment when compared to a vehicle (VH, in blue) treatment in both the sham groups (n=10) and the traumatic brain injury (TBI) groups (n=29-30). This pattern persisted into the 7-week treatment for both the sham (n=10) and TBI (n=19) groups under VH or PP treatments. Statistical significance was assessed using Bonferroni’s multiple comparisons test (*p<0.033, **p<0.002, ***p<0.001). (e-g) Principal Coordinate Analysis (PCoA) ordination plots were based on Weighted UniFrac distances and PERMANOVA assessed group dissimilarities. Beta diversity did not show significant differences between the VH and PP groups after 2 weeks of treatment nor between sham groups after 7 weeks of treatment. However, a significant variation in beta diversity was observed between VH and PP groups in TBI mice at the 7-week treatment (g). *Firmicutes/Bacteroidetes* ratio decreased in the VH group after two weeks compared to the VH from sham mice after 7 weeks of treatment in males (h) and females (i). In addition, the TBI group had an increased ration compared to the Sham mice at 7 weeks of treatment. No differences were observed in this ratio within the PP groups. Statistical significance was assessed using Bonferroni’s multiple comparisons test (*p<0.033, **p<0.002, ***p<0.001).

The temporal dynamics of microbiota genera following VH and PP treatments in mice revealed sex-dependent modifications in the microbiota composition. At 2-week of treatment, the relative abundance of the top 20 microbial genera in mice treated with VH exhibited a striking sex-dependent pattern. In male mice, there was a pronounced presence of *Dubosiella*, contrasting with the female mice, who showed a higher abundance of *Faecalibaculum*, particularly within those subjected to TBI-VH treatment (**Fig. 4 a**). In the PP-treated groups, sham male mice demonstrated a higher frequency of *Lactobacillus*, while an increased presence of Faecalibaculum characterized their female counterparts. In TBI-PP mice, a notable dominance of *Dubosiella* was observed in males, many of which also harbored *Faecalibaculum*. In contrast, female mice predominantly displayed an abundance of *Faecalibaculum* (**Fig. 4 b**). *Lactobacillus* was identified as the most prevalent genus throughout these subsets, indicating its significant role in the gut microbiota composition under these experimental conditions. By 7-week of treatment there was a notable balance in the microbial composition between male and female VH-treated mice, regardless of whether they were subjected to sham or TBI interventions (**Fig. 4 c**). Conversely, in the PP-treated groups, sex-dependent variations were more pronounced. Sham-treated male mice predominantly hosted *Ileibacterium*, a genus not previously highlighted at an earlier time point. TBI-exposed female mice, however, showed a higher abundance of *Lactobacillus* relative to their male counterparts, underscoring the ongoing sex-specific influence of PP treatment in the context of TBI (**Fig. 4 d**).

**Figure 4.**
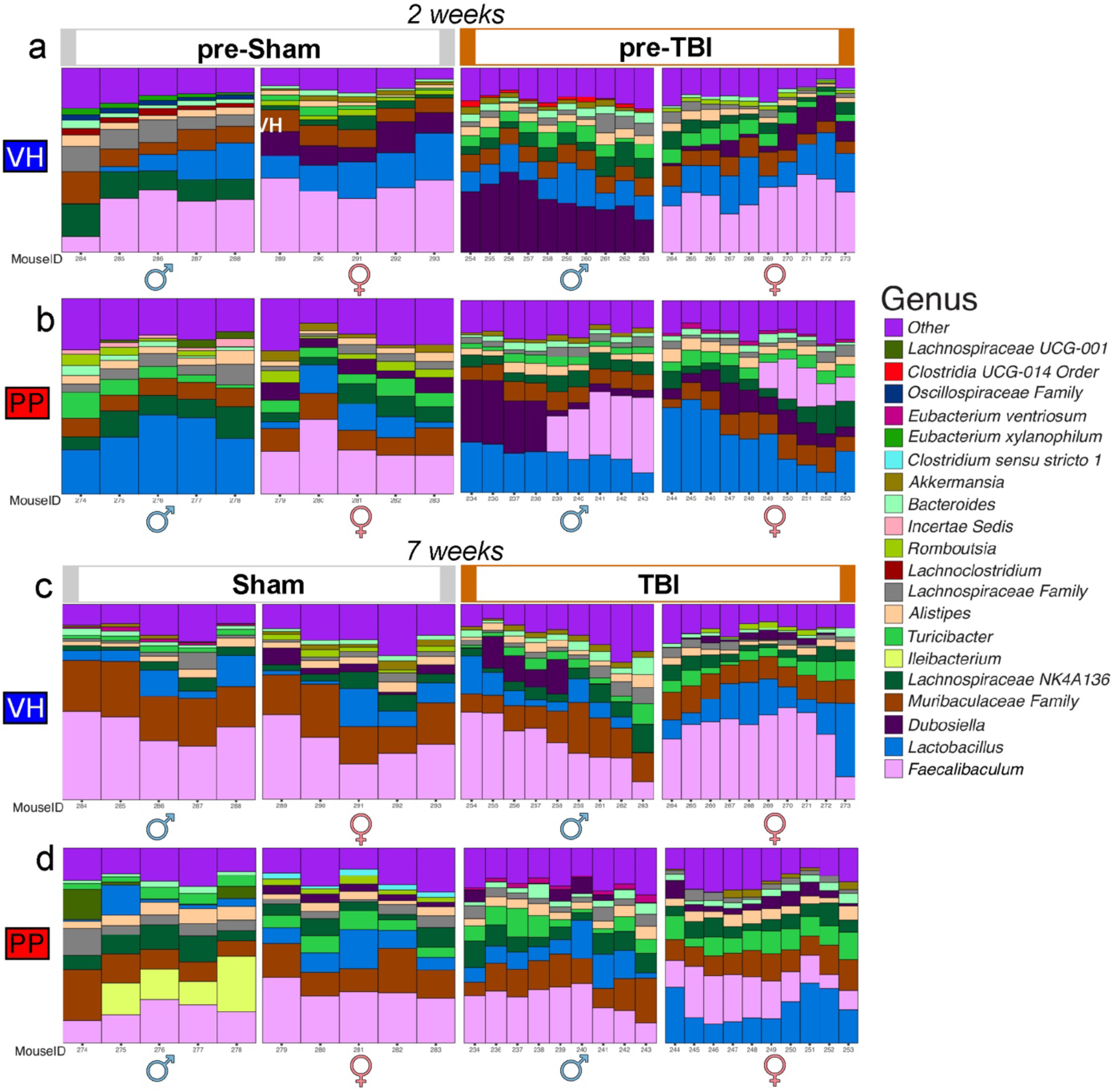
Temporal dynamics of microglia genera in VH and PP-treated mice is sex-dependent. (a-b) At the 2-weeks, the relative abundance of the top 20 microbial genera in VH-treated mice pre-sham and pre-TBI, noting the pronounced presence of *Dubosiella* in males compared to females, who exhibit a higher abundance of *Faecalibaculum* within the TBI-VH subset. (b) Sham-PP male mice are characterized by a higher frequency of *Lactobacillus* and females by *Faecalibaculum*. TBI-PP group reveals a dominance of *Dubosiella* in males, with a significant portion also harboring *Faecalibaculum*, whereas females predominantly present with *Faecalibaculum*. Across both subsets, *Lactobacillus* is the most prevalent genus. (c-d) Delineating the microbial landscape at 7-weeks, (c) a balanced microbial composition between males and females in VH-treated mice, irrespective of sham or TBI. Contrastingly, in the PP group, sham-treated males predominantly host *Ileibacterium*, whereas TBI-exposed females exhibit a higher abundance of *Lactobacillus* relative to males. Each stacked bar represents an individual mouse’s gut microbiota composition, with unique colors corresponding to different genera as indicated in the legend to the right of the panels. The bars show the variation within groups, highlighting the dynamics of the fecal microbiome in response to sham and TBI treatments over time. Unclassified Taxa were excluded. Statistical significance was again assessed using Bonferroni’s multiple comparisons test (*p<0.033, **p<0.002, ***p<0.001).

The comparative analysis focused on the microbial abundance variations observed between the 2-week and 7-week intervals in mice subjected to VH and PP treatment across both sham and TBI conditions (**Fig. 5**). In the sham VH-treated groups, both male and female mice exhibited no significant alterations in microbial taxa after 2 weeks of treatment (**Fig. 5 a, b**), suggesting resilience of the gut microbiota to the sham procedure without notable disruptions. We observed a shift in the PP male group with the sham mice leading to significantly increased abundance of *Lactobacillus gasseri (*LFC=1.91, q=0.004), *Lactiplantibacillus* genera (LFC=2.53, q=0.009), *Lactobacillus* genera (LFC=2.10, q=0.027), and *Limosilactobacillus* genera (LFC=1.91, q=0.029) (**Fig. 5 c**). A disparity was seen between the males and the females of the sham-PP groups as only the *Lactiplantibacillus* genera (LFC= 2.89, q= 0.043) was shown to increase in abundance within the PP female cohort statistically. Investigating the effect of TBI in both males and females, VH-treated mice did not demonstrate any significant shifts in microbial taxa, underscoring a consistent microbial profile (**Fig. 5 e, f**). The analysis encompassing PP-treated mice in the TBI scenarios (**Fig. 5 g, h**) was drastically altered compared to the sham group. TBI-PP treatment of male mice was shown to significantly increase *Lactiplantibacillus* (LFC=1.96, q= 8.64e-5) and *Lactobacillaceae_HT002* genera (LFC= 2.44, q=3.10e-4). In parallel, a significant reduction in the species *Lactobacillus helveticus* (LFC= -1.84, q=5.90e-4) was also seen in the TBI-PP-treated male mice. The TBI-PP-treated female mice (**Fig. 5 h**) underwent the most significant microbial changes with increases in the abundance of *Lactiplantibacillus* genera (LFC= 2.41, q= 4.32e-7), *Faecalibaculum genera* (LFC= 2.90, q=0.002), *Lactobacillaceae_HT002* genera (LFC=1.98, q=1.99e-4), *Limosilactobacillus* genera (LFC=1.42, q=1.94e-3), *Lactobacillus_gasseri* (LFC=1.20, q=0.002), and *Lachnospiraceae NK4A136* genera (LFC=1.72, q=0.013).

**Figure 5.**
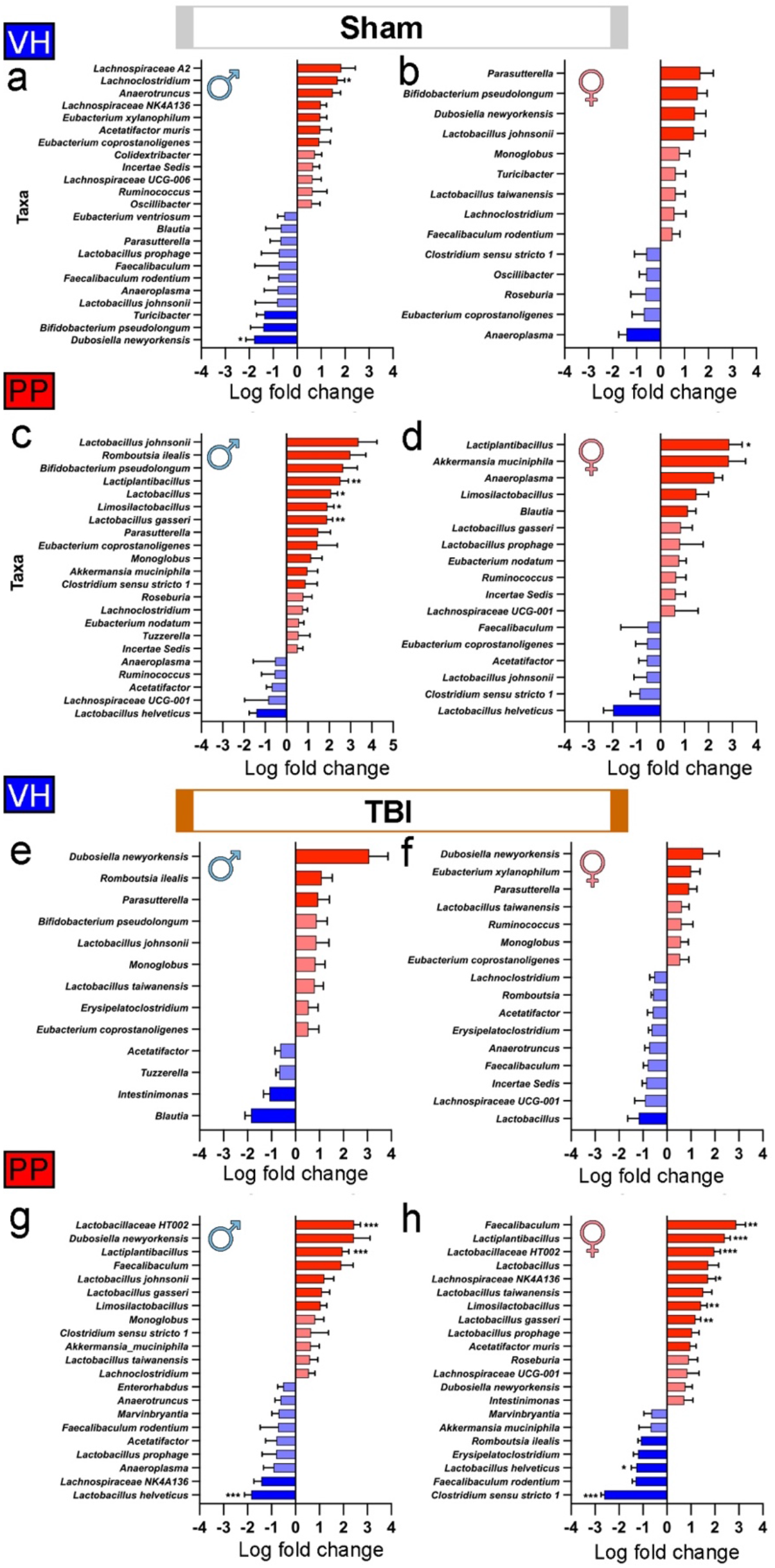
Differential abundance of microbial taxa in VH and PP groups under sham and TBI conditions. The shifts in microbial communities following sham and TBI interventions between the 2-week and 7-week time frames for mice treated with the VH in both sham (a-d) andTBI (e-h) groups, as well as for VH and PP treated mice. The mean log fold change in abundance of a particular taxon, with the blue bars indicating a decrease and red bars indicating an increase in abundance after 7-weeks of treatment compared to the 2 weeks of treatment. Taxa names are listed on the y-axis, and the log fold change is plotted on the x-axis. In the sham treated with VH, both male (a) and female (b) groups showed no notable differences in microbial taxa. In the sham mice treated with PP, males (c) exhibited a significant rise in the abundance of *Lactiplantibacillus, Lactobacillus, Limosilactobacillus,* and *L. gasseri*. Females in the sham-PP group significantly increased, specifically in *Lactiplantibacillus* (d). In the TBI mice, neither males (e) nor females (f) in the VH group demonstrated significant changes in taxa. However, males (g) in the TBI-PP group presented an increased abundance of *Lactobacillaceae* and *Lactiplantibacillus*, alongside decreased *L. helveticus*. Females (h) in the TBI-PP group revealed taxonomic alterations characterized by increased levels of *Faecalibaculum, Lactiplantibacillus, Lactobacillaceae, Lachnospiraceae,* and *Limosilactobacillus*, as well as *L. gasseri,* with a concurrent reduction in the abundance of the taxa *L. helveticus* and *Clostridium*. Error bars denote the standard error of the mean (SEM). A log-fold-change minimum threshold of 0.5 was used within the ANCOMBC2 visualization. Statistical significance was determined using the resulting q-value from the ANCOMBC2 analysis with (*q<0.05, **q<0.01, ***q<0.001).

Equivalently, some taxa significantly decreased in the female TBI-PP-treated mice, including *Clostridium sensu stricto 1* (LFC= -2.61, q=1.18e-4) and *Lactobacillus helveticus* (LFC= -1.27, q= 0.017) (**Fig. 5 h**). We also modeled microbial abundances using beta-binomial regression [50]. This yielded concordance with the ANCOM results, identifying significant shifts in taxonomic profiles in sham-VH and TBI-VH males compared to sham-PP and TBI-PP in males and TBI-PP in females (**Supplemental Fig. 1**).

### Germ-free mice show a higher lesion level than wild-type mice after TBI, and pan-probiotics treatment reduces the lesion and cell death following TBI in males

GF male mice showed around 33% more lesion volume than wild-type male mice at 3 dpi (****p<0.0001) (**Fig. 6 a**), indicating the critical role of microbiota in brain damage repair. In our examination of the effects of a PP treatment on TBI, we observed that lesion volume showed a significant reduction in male subjects treated with PP compared to VH (*p < 0.05). At the same time, females did not exhibit a significant change (**Fig. 6 b**). Representative cresyl-violet brain sections highlight the areas of the lesion (**Fig. 6 c**). TUNEL assay results indicated a significant decrease in dying cells in males receiving PP treatment compared to VH (*p < 0.05), with females showing a less pronounced lesion volume compared to male counterparts in the VH group (*p < 0.05) (**Fig. 6 d, d1-4**). These findings suggest that the PP treatment may confer neuroprotective effects in a sex-specific manner post-TBI.

**Figure 6.**
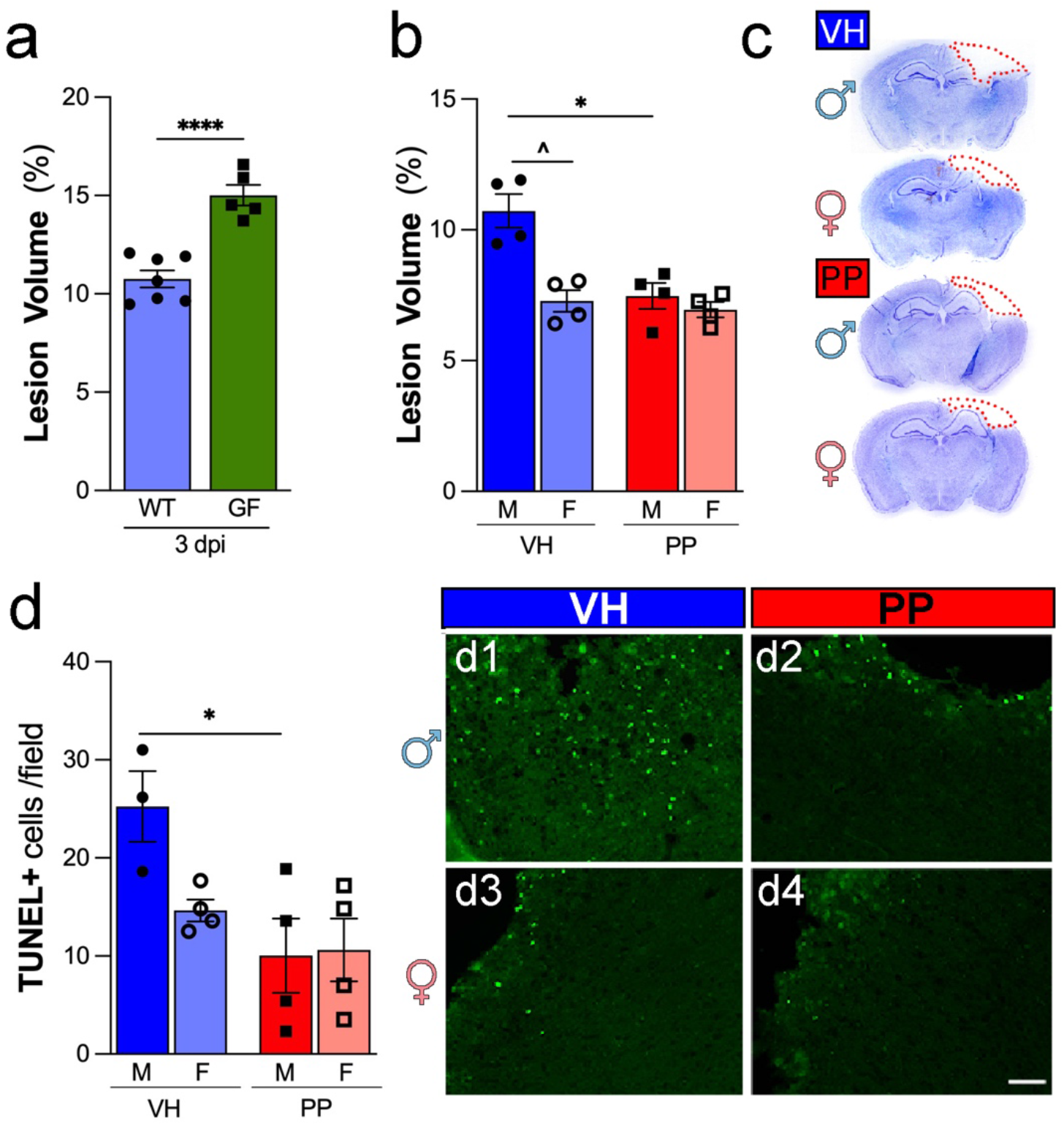
Pan-Probiotic Treatment decreases Lesion and Neuronal Damage. (a) Germ-free (GF) male mice showed around 33% more lesion than wild-type male mice at 3 days post-injury (dpi). (b) Male mice subjected to traumatic brain injury (TBI) exhibit significantly reduced mean lesion volumes after pan-probiotic (PP) treatment in comparison to the vehicle-treated (VH) group. (c) Brain sections stained with cresyl-violet, featuring delineating the lesion area comprising the cavity and edematous region at 3 dpi. (d) Quantitative analysis demonstrates a substantial decrease in the number of TUNEL + cells per field in the injured cortex of PP-treated mice as opposed to VH-treated counterparts. (d1-4) Representative images displaying TUNEL + cells in the cerebral cortex following TBI providing visual confirmation of the quantified data. The data represents n=3-4 mice per group, with significance determined through one-way ANOVA with post-hoc Tukey’s multiple comparison test (*p<0.05, **p<0.01, ***p<0.001, ****p<0.0001). Scale bars represent 50 µm in d1-4.

### Administering pan-probiotics diminishes the activation of microglia/macrophages in males three days following traumatic brain injury

To evaluate the effects of PP treatment on the anti-inflammatory activity after acute TBI, we analyzed the activation of the microglia and macrophage population using the Iba-1 marker. The quantitative analysis of Iba-1+ cells revealed a significant decrease in cell density in male and female mice treated with PP compared to the VH group 3 dpi (**Fig. 7 a, c-f**). The Iba-1+ area within the observed fields also showed a decrease, with the male PP group demonstrating a significant enlargement in comparison to the male VH group (***p<0.001) (**Fig. 7 a, c-f**). The Iba-1+ area within the observed fields also showed an increase, with the male VH group demonstrating a significant enlargement in comparison to the male PP group (**p<0.01) (**Fig. 7 b, c-f**). High-magnification images of Iba-1+ cells (**Fig. 7 j-m**) highlighted the morphological differences, with the VH-treated microglia appearing to have a more activated phenotype. The Sholl analysis was performed, and data indicated a reduced complexity in the dendritic architecture in PP-treated males (**Fig. 7 g-I)**, as shown by the decreased number of intersections (**Fig. 7 n**) and the shorter average length of processes (**Fig. 7 p**) at distances greater than 10 µm from the cell body compared to the VH-treated mice. In addition, F4/80+ macrophages analysis revealed a significant elevation in cell density in VH-treated males in comparison to PP male mice (**p<0.01) (**Fig. 7 r**), aligning with the observed alterations in microglia morphology and suggesting a potential modulatory effect of PP on microglia activation. The females were not statistically significant when comparing PP and VH treatments (**Fig. 7 o, q**). Representative images of F4/80+ staining corroborate the quantified data, with a noticeable reduction in signal intensity in the PP-treated groups compared to the VH-treated groups (**Fig. 7 s-w**). Overall, the findings suggest that post-TBI microglial activation and morphological changes are reduced by PP treatment, with more pronounced effects observed in males.

**Figure 7.**
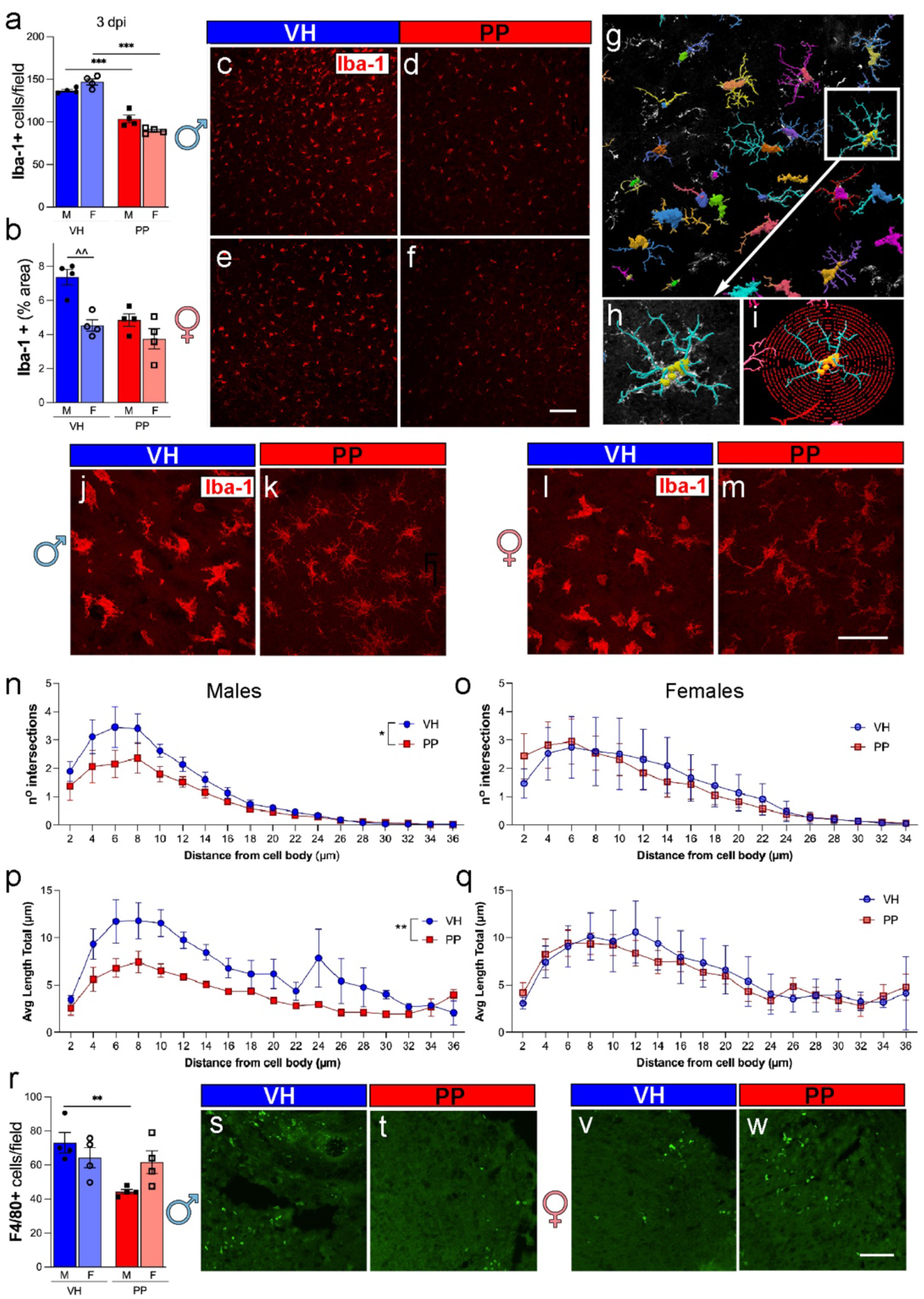
Pan-Probiotic Treatment decreases Microglia/Macrophage Activation. (a) Quantitative analysis shows a significant reduction in the number of microglia/macrophages (Iba-1+ cells) in mice subjected to Pan-Probiotic (PP) treatment compared to those treated with the vehicle (VH) as well as a significant decrease in PP-treated female mice compared to PP-treated males at 3 days post-injury (dpi). (b) The percentage of the area occupied by Iba-1+ cells also exhibits a noteworthy decrease in PP-treated male mice compared to their VH-treated counterparts and the VH-treated females compared with VH-treated males (c-f) Representative images depict Iba-1+ cells in the injured cortex of mice treated with VH and PP. (g) A representative image of Neurolucida 360 tracing software illustrates the presence of Iba-1+ cells. (h) A zoomed-in representative image highlights traced microglia. (i) An image displays concentric circles (red) drawn at a 2 μm distance for Sholl analysis. (j-m) Representative confocal microscopy images of Iba-1+ staining in PP and VH brains at 40X magnification. The Sholl analysis reveals significant differences in the number of intersections from the distance of the cell body of Iba-1+ cells (n) and the total average length of the branch from the distance of the cell body (p) in the injured cortex of males treated with PP compared to VH. No differences were observed in female mice (o, q). (r) Quantitative analysis demonstrates a significant reduction in the number of macrophages (F4/80+ cells) in the injured cortex of males treated with PP compared to VH-treated males, with no such effect observed in females. (s-w) Representative images illustrate F4/80+ cells in the injured cortex of PP- and VH-treated male and female mice. The data represents n=4 mice per group, and statistical significance was determined using one-way ANOVA with post-hoc Tukey’s multiple comparison test. For the Sholl analysis, the area under the curve was calculated, and an unpaired t-test using the mean, SEM, and n was utilized (*p<0.05, **p<0.01, ***p<0.001). Scale bars represent 50 µm in c-f and 20 µm in j-m, s, and w.

### Pan-probiotic treatment improves motor and partiality and reduces anxiety-related behaviors associated with TBI

In assessing behavioral outcomes following TBI, male and female mice exhibited distinct responses after PP treatment across various tests. In the rotarod test, males demonstrated a significant decrease in latency to fall post-TBI compared to sham-operated controls, indicating a motor deficit as previously established in our mouse CCI model. This effect was most pronounced in TBI-VH treated males at early time points (1, 3, and 7 dpi, ###p<0.001), with gradual recovery over 35 dpi, yet still significantly lower than sham-VH groups (##p<0.01 at 14 dpi and not significantly different at 35 dpi). Females showed a similar motor impairment, with TBI-VH groups exhibiting reduced latency to fall compared to sham-VH groups (###p<0.001 at 1 and 3 dpi, ##p<0.01 at 7 dpi) (**Fig. 8 a, b**). PP treatment significantly improved the motor deficits induced by TBI in both male and female mice (**Fig. 8 a, b)** (*p<0.05 at 1 and 3 dpi for males and at 1 and 7 dpi for females). In the EPM, both male and female sham-PP and TBI-PP groups spent more time in the open arms entries, showing a non-significant trend towards increased compared to VH groups (p=0.11 and p=0.06, respectively) (**Fig. 8 c**).

**Figure 8.**
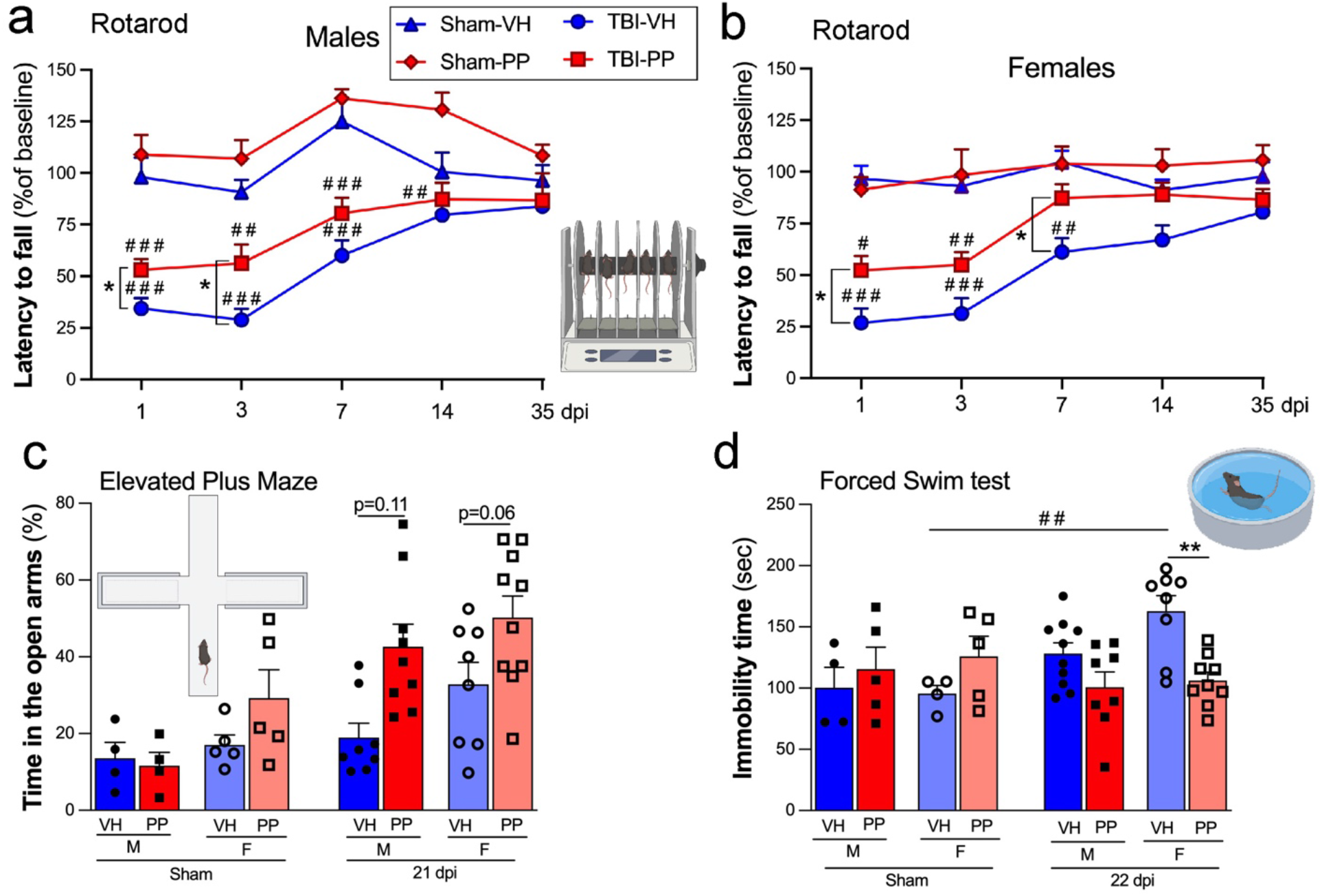
Pan-Probiotic Therapy Enhances Motor Skills and Reduces Anxiety/Depression-Related Behaviors After Traumatic Brain Injury (TBI) in a Sex-Dependent Manner. Motor behavior was evaluated using the Rotarod test for both male (a) and female (b) mice, which were administered either pan-probiotic (PP, in red) or a Vehicle water solution (VH, in blue). For males (a), notable differences were observed on days 1, 3, and 7 days between Sham and TBI mice receiving VH on days 1, 3, 7, and 14 between sham and TBI mice treated with PP. Specifically, on days 1 and 3, a significant disparity was seen between VH and PP treatments in the TBI group. In females (b), notable differences were observed on days 1, 3, and 7 between sham and TBI mice receiving VH and on days 1 and 3 days between Sham and TBI mice treated with PP. Also, differences were seen on days 1 and 7 between the treatments of VH and PP in the TBI groups. (c) In the Elevated Plus Maze test conducted 21 days after TBI, TBI-PP-treated female mice exhibited a significant increase in the number of entries into open arms compared to sham-VH females, and similarly, TBI-PP-treated males showed more entries compared to their sham-VH counterparts. In the Forced Swim Test (d), while male mice showed no significant differences, female mice treated with PP demonstrated a notably longer duration of immobility, 22 days post-TBI, compared to those treated with VH. Sham vs. TBI (## p<0.01, ### p<0.001, #### p<0.0001), and VH versus PP (*p<0.05, **p<0.01).

The FST revealed a significant reduction in immobility time observed between females sham-VH and TBI-VH groups (##p<0.01). Additionally, female TBI-PP mice spent significantly less time immobile than TBI-VH groups (**p<0.01), suggesting an effect of the PP treatment in improving the depressive-like behavior induced by TBI. In contrast, male TBI-PP subjects did not show significant differences in immobility time compared to TBI-VH groups, although a trend toward decreased immobility was noted (**Fig. 8 d**). Sham-operated controls did not exhibit any significant changes across treatments.

### Pan-probiotics did not alter the gut permeability and systemic inflammation

In a histological examination of the intestines following TBI, both male and female subjects displayed changes in gut morphology at different time points post-injury. At 3 dpi, villi length in the VH-treated males and females appeared similar to sham controls; however, PP treatment resulted in a reduction in villi length in males that was not observed in females (**Fig. 9 a, a1-4, b**). By 35 dpi, villi length in the PP-treated males significantly recovered. In contrast, PP-treated females showed a nonsignificant trend towards increased villi length compared to VH-treated counterparts (**p<0.01) (**Fig. 9 c, c1-4, d**). Mucin+ area analysis revealed no significant differences between the groups at 3 dpi (**Fig. 9 e, e1-4, f**), and mucin+ goblet cell count did not differ significantly (**Fig. 9 g).** However, at 35 dpi, mucin+ goblet cell numbers in PP-treated males were slightly increased compared to VH-treated males, though not reaching statistical significance (**Fig. 9 h, h1-4, i, j**). As a systemic inflammatory marker, SAA levels showed no significant differences between groups. However, the PP treatment maintains a trend of reduction in SAA levels compared to the VH groups, suggesting that PP treatment reduces the acute phase response at 3 dpi, mainly in males (**Fig. 9 k**). Additionally, body weight measurements for 35 days indicated no significant differences between groups, with all mice showing a typical growth curve regardless of sex or injury (**Fig. 9 l**). Collectively, these results suggest that while PP treatment reduced villi length in females, the overall absence of significant changes in mucin production, goblet cell number, systemic inflammation (as measured by SAA levels), or body weight suggests that the observed histological alterations in the gut may be localized responses to TBI and treatment, rather than systemic effects.

**Figure 9.**
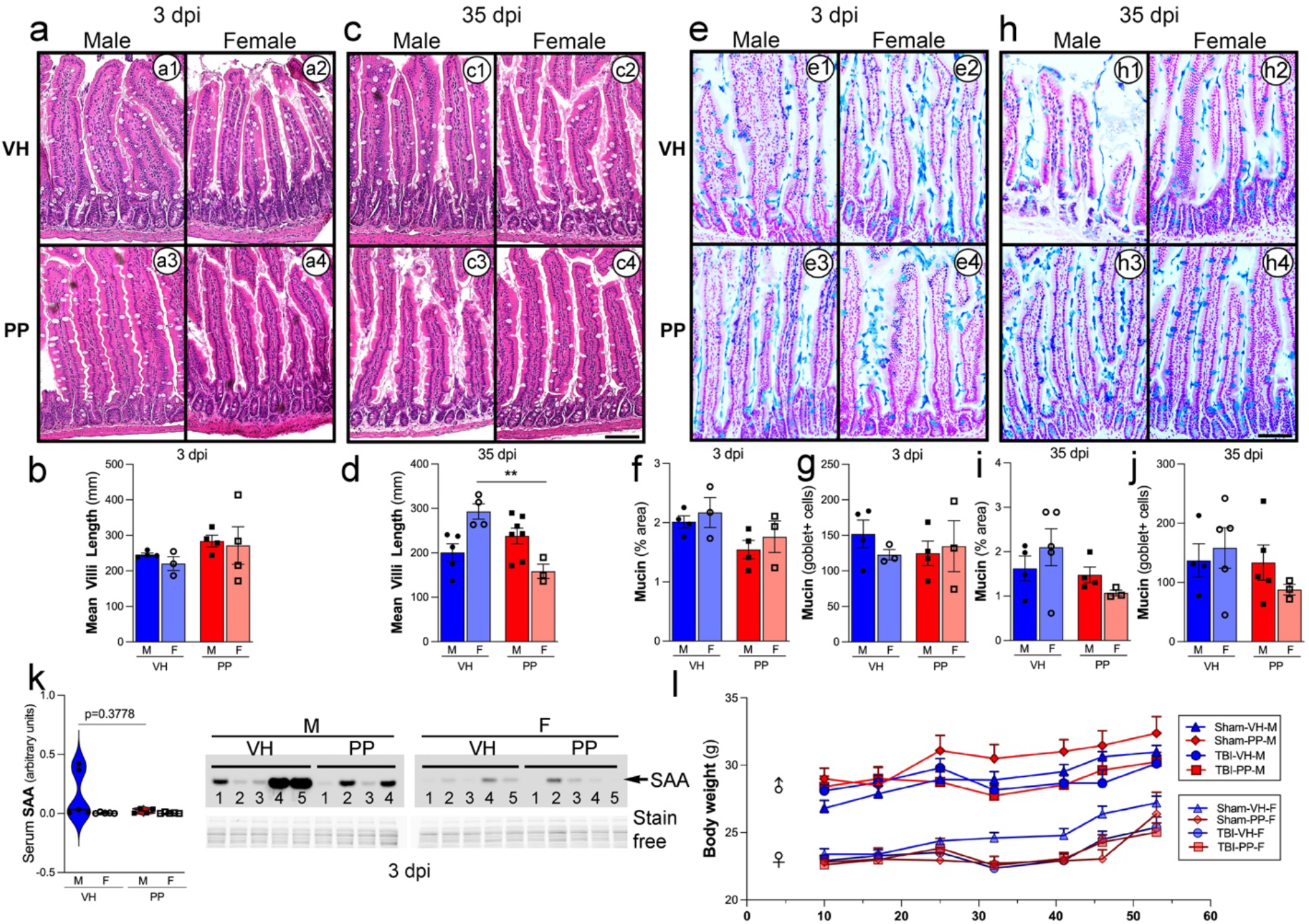
Probiotic treatment restored gut morphology and slightly enhanced mucin production after traumatic brain injury (TBI) in male mice. Histological assessment of the intestinal morphology and mucin production was conducted at 3- and 35-days post-injury (dpi) in both male and female mice, using either a vehicle (VH) or a pan-probiotic treatment (PP). At 3 days post-TBI, there was no significant change in the length of the villi in the colon across all groups (a, a1-4, b). At 35 dpi, male PP-treated mice showed a significant increase in villus length compared to their VH counterparts, highlighting the PP restorative potential (c, c1-4, d). When assessing mucin production in the colon, no significant differences were noted at 3 dpi regarding the mucin+ area or the count of mucin+ goblet cells across any group (e, f). The number of mucin+ goblet cells followed a similar trend (g). At 35 dpi, although not statistically significant, an increase in both mucin area and goblet cell numbers in the PP-treated groups hinted at a delayed beneficial effect of the probiotics (h, i). Serum amyloid A (SAA) levels, a marker of inflammation, remained unchanged across treatments and sexes. However, the PP group tends to show a more uniform reduction in SAA levels than VH (k). The study also observed body weight changes, indicating that TBI influenced weight gain. PP-treated mice showed a trend towards quicker weight recovery than those given VH across both sexes (j). VH versus PP (**p<0.01). Scale bars (a1-4, c1-4, e1-4, h1-4) represent 50 µm.

## DISCUSSION

In recent years, research has demonstrated the pivotal role of the gut-microbiome-brain axis in maintaining neurological health [51]. The gut microbiota can affect CNS functions in both healthy physiological conditions, encompassing neurotransmission, neurogenesis, and glial activation [52]. In the context of neuroinflammatory diseases, there is a significant alteration in the composition and diversity of the microbiome, which influences the progression of neuropathological conditions [53]. This phenomenon is observed in individuals who have suffered brain damage, who demonstrate increased systemic inflammation, changes in the composition of gut microbiota, disrupted intestinal motility, and neuroinflammation [54–56].

The temporal dynamics of microbial communities elucidated in this study provide valuable insights into the sex-dependent effects of probiotic treatment in the context of brain injury. Our study showed that gut microbiota dysbiosis following TBI can be reversed by administering PP, which concurrently exhibits neuroprotective effects within the injured brain. Our previous work showed the impact of TBI on bacterial dysbiosis corresponding to relative abundance and the diversity of the *Lactobacillaceae* family [14], which is one of the ubiquitously seen healthy bacteria in the human gut. *Lactobacillus*, a genus of bacteria belonging to the phylum *Firmicutes* [57], have been shown to have neuroprotective effects [58]. A probiotic regimen consisting of *Bifidobacterium, Lactobacillus,* and *Enterococcus faecalis* markedly reduced levels of pro-inflammatory cytokines post-treatment in severe TBI patients [59]. In our study, we observed a dramatic decrease in *Lactobacillus gasseri* at the species level at 24 h post-TBI [14]. We hypothesized that the administration of a probiotic mixture consisting of various *Lactobacilli* strains, specifically *L. plantarum, L. reuteri, L. helveticus, L. fermentum, L. rhamnosus, L. gasseri, and L. casei.* might have a neuroprotective action during the acute and chronic phases in a mouse model of TBI. Our selection was supported by literature documenting the beneficial effects of the *Lactobacillus* strains in various animal model studies, where modulation of the inflammatory response was a fundamental aspect of the disease process [60–63]. Different strains of *Lactobacillus* have been shown to decrease the production of inflammatory cytokines such as TNF-α and IL-6 [64, 65], downregulate pro-apoptotic neurons, and activate inflammatory mediators [66]. The administration of *Lactobacillus* has also been shown to mitigate locomotor dysfunction, anxiety, and depression after brain insults [67, 68]. The probiotic *L. acidophilus* has been shown to help repair the intestinal mucosal barrier and improve the terminal ileum villus morphology and GI motility [69], and also enhances the microbial community’s diversity [70]. While the administration of probiotics does not correlate with their ability to colonize the gut permanently, research has shown that even if probiotics do not establish long-term residence in the intestines, their introduction can still modify the existing intestinal microbiota composition [71].

The positive impacts of probiotics are driven by their metabolite profiles, like SCFAs, regardless of whether the probiotic strain establishes long-term colonization in the intestine [66, 71]. SCFAs are recognized for their neuroprotective [72], neuronal survival, plasticity [73], and anti-inflammatory properties [24]. They also modulate microglial activation [74], maturation and function [75] across various conditions [76]. *Lactobacilli* produce SCFAs such as acetate, propionate, and butyrate [77]. Our findings indicate that PP treatment for 7 weeks increases the relative abundance of SCFAs in serum in a favorable metabolic environment for TBI recovery. Interestingly, SCFA levels did not show significant changes at 3 dpi. This might indicate that the acute neuroinflammatory response to TBI abolishes the production of SCFAs by selected PP. However, PP administration increases SCFA levels once the inflammation is restored, evidenced at 35 dpi. We have not observed differences between male and female mice, yet other studies have documented sex-based differences in SCFA profiles [78, 79]. These variations are likely attributable to the temporal course of hormonal disparities following TBI [29].

There were increases in microbial counts and significant variations in beta diversity, especially pronounced in TBI mice for 7 weeks of PP administration. Alpha diversity analysis also indicates an augmentation in the sham-treated male mice receiving PP compared to those treated with VH, with no similar increase observed in females, underscoring the gender-specific microbial response. Our results suggest that the PP administration corresponded with changes in phyla like *Firmicutes* and *Bacteroidetes* and alterations in the gut microbiota that were specific to each sex. Our findings suggest a significant increase in the abundance of genera such as *Lactiplantibacillus, Lactobacillus,* and *Limosilactobacillus*, as well as the species *L. gasseri* in male mice, which could be indicative of the PP targeted effect on male microbiota. We found specific shifts in microbial taxa in the PP treated mice, such as the increased abundance of beneficial bacteria like *Lactobacillaceae*, *Limnosilactobacillus* and *Lactiplantibacillus* populations, but a decline in *L. helveticus,* suggest that PP treatment could correct dysbiosis associated with TBI. Previous reports associated *Limnosilactobacillus* and *Lactiplantibacillus* with anti-inflammatory properties and gut health [80]. Females in the TBI-PP group exhibited more complex taxonomic changes post-TBI with notable increases in taxa such as *Faecalibaculum, Lactiplantibacillus, Lactobacillaceae,* and *Lachnospiraceae*, along with *Limosilactobacillus* and *L. gasseri*. The increase of *Clostridium* taxa could suggest a nuanced interaction between the administered probiotic and host physiology in the injury context [81]. We found an increase in *Faecalibaculum* in the TBI-PP-treated group. This bacterium is likely associated with the observed reduction in depression and anxiety [82]. Surprisingly, the decline of *L. helveticus* in both sham and TBI-PP mice may reflect a rebalancing of the gut ecosystem, possibly correcting the associated dysbiosis providing an opportunity for other bacteria to compete more effectively for resources than *L. helveticus*. Also, different strains may produce substances that inhibit the growth of potentially beneficial bacteria [83].

A limitation of our current study is that the characterization of gut dysbiosis through short-read 16S sequencing is constrained by the small amplicon size, leading to possible taxonomic misidentification [84]. 16S amplicon sequencing is much more affordable and scalable than shotgun metagenomics. However, the inability to confidently classify bacteria at the species and strain levels and the lack of accurate functional annotations that can be performed are major detriments when attempting to understand the gut microbiome changes [85]. One promising area that spans the gap between short-read amplicon sequencing and shotgun metagenomics is nanopore full-length amplicon sequencing, which has been gaining popularity in microbiome research through its ability to identify bacteria taxa at the species level [86] which we will implement in future studies.

Our study observed a tendency for PP treatment to diminish the levels of SAA in serum, a protein associated with acute-phase peripheral inflammation, aligning with documented evidence that probiotic supplementation can lessen the occurrence of infections [87]. Our findings indicate no changes in the intestinal epithelium, implying that the PP-administered treatment does not adversely influence gut integrity. Concerning neuroprotective outcomes, our data indicates that PP exerts a sex-specific impact, predominantly evidenced by decreased lesion volume and cell death in male mice aligning with the observed neuroprotective outcomes, a finding that is consistent with outcomes reported in our prior research [29]. Probiotic supplementation has been shown to mitigate apoptosis and enhance inflammatory states in animal models. For instance, the administration of *C. butyricum* diminished neuronal cell death triggered by bilateral common carotid artery occlusion in diabetic mice [88]. Another investigation using broiler chickens as the model found that *L. plantarum* supplementation decreased apoptosis and intestinal inflammation, an outcome attributed to enhanced SCFA production and alterations in the microbiota composition [89]. Our results show how animals lacking microbiota, GF mice, respond to TBI with extensive brain injury. Previous studies have indicated that supplementing GF mice with SCFAs ameliorates abnormalities in microglia [75].

The host-microbiota plays a role in microglial maturation and activation [90], and as the resident immune cells of the CNS, play a pivotal role in the brain’s response to injury, shifting from a ’resting’ state to an ’activated’ state [91], and releasing proinflammatory cytokines [4]. Interestingly, our findings indicate that PP treatment exerts a differential effect on the microglial activation state, as evidenced by a significant decrease in Iba-1+ cell density and alterations in cell morphology, particularly in male mice. Moreover, the observed reduction in F4/80+ macrophages in male mice receiving PP treatment may also influence peripheral immune cells infiltrating the brain after injury. The more marked effects seen in male, relative to female mice, align with an increasing body of evidence suggesting that male and female brains may exhibit distinct responses to TBI and subsequent treatments, a phenomenon we have previously demonstrated [29, 92, 93]. *Lactobacillus* supplementation has been linked to reductions in anxiety and depression [67], [68]. In our study, PP treatment tends to mitigate anxiety-related behaviors, as evidenced by the performance in the EPM and the FST in males, and an improvement in motor functions. A study in neonatal mice shows how infection with *C. rodentium* determined the role of *L. rhamnosus* and *L. helveticus* in mitigating corticosterone-induced colitis and stress-induced cognitive dysfunctions [94]. Moreover, a novel composite probiotic mixture (VSL#3) containing diverse strains of *Bifidobacteria* and *Lactobacilli* prevented the diet-induced memory deficits, however, no differences were observed in anxiety-like behavior [95]. In a clinical study, participants who received a 4-week multispecies probiotics intervention showed a significantly reduced overall cognitive reactivity to sad mood [96]. Consequently, in our research, employing PP appears to ameliorate the anxiety and depressive behaviors triggered by TBI, a finding that aligns with trends observed in other animal studies.

## CONCLUSIONS

In conclusion, our study provides compelling evidence of the potential benefits of PP supplementation in TBI. The modulation of the gut microbiome, metabolic profiles, and inflammatory processes, coupled with the neuroprotective effects and behavioral improvements observed, underscore the therapeutic promise of probiotics in a sex-dependent manner. The insights provided by these findings emphasize the potential of targeting the gut microbiota and its metabolic pathways as a novel therapeutic strategy for TBI. These outcomes provide a promising avenue for developing probiotic-based interventions in TBI. However, further research is needed to fully elucidate the mechanisms underlying these effects and their long-term implications. These results collectively contribute to a burgeoning discourse on the gut-brain axis, emphasizing the potential of probiotic treatments in modulating the gut microbiome post-TBI, with implications for therapeutic interventions and recovery.

## ABBREVIATIONS

AhR: aryl hydrocarbon receptor
ANOVA: Analysis of variance
ASVs: amplicon sequencing variants
BBB: Blood-brain barrier
BCL: Binary base call
BDNF: brain-derived neurotrophic factor
CCI: Controlled cortical impact
CFU: Colony forming unit
CNS: Central nervous system
CRP: C-reactive protein
dpi: Days post-injury
EPM: Elevated Plus Maze Test
FST: Forced Swimming Test
GF: Germ-Free
HEPA: High-efficiency particulate air
MFI: Mean fluorescence intensity
MRM: Multiple reaction monitoring
NGS: Normal goat serum
PBS: Phosphate-buffered saline
PBS-T: PBS enhanced with 0.5% Triton X-100
PBS-Tw: PBS-Tween 20
PCR: Polymerase chain reaction
PP: Pan-Probiotic
rpm: Revolutions per minute
SAA: Serum amyloid A
SCFAs: Short-chain fatty acids
TBI: Traumatic brain injury
TUNEL: Terminal deoxynucleotidyl transferase dUTP nick end labeling
VH: Vehicle

## ACKNOWLEDGEMENTS

The authors thank the CPRIT Proteomics and Metabolomics Core Facility (RP210227), NIH (P30 CA125123), and Dan L. Duncan Cancer Center at Baylor College of Medicine. We also thank the Pathology and Histology Core at Houston Methodist Research Institute the Baylor College of Medicine Gnotobiotic Rodent Facility, and the Center for Comparative Medicine, especially Alton G. Swennes, for his help with the germ-free mice. The authors thank Cathryn Gunawan, Eric Wang, Manoj Dandekar, Shashi Krishnamurthy, and Pranjal Sheth for their technical help. The authors also thank Anna Dodson for editing the text and for the philanthropic funding from Paula and Rusty Walter and the Walter Oil & Gas Corp Endowment at Houston Methodist. Figure 1 was done utilizing Biorender.com.

## AUTHOR CONTRIBUTIONS

M.H., H.F., S.S., A.M., T.T., and S.V. initiated, designed, planned, and oversaw all aspects of the study. M.H., H.F., S.S., A.M., and S.V. performed the experimental work and data analysis and drafted the manuscript. All authors reviewed and edited the final version of the manuscript.

## FUNDING

This work was supported by Houston Methodist Research Institute, NIH grant R21NS106640 (SV) from the National Institute for Neurological Disorders and Stroke (NINDS), and NIH grant R56AG080920 (SV) from the National Institute on Aging (NIA).

## AVAILABILITY OF DATA AND MATERIALS

The data used in the study are available from the corresponding author upon reasonable request. Sequence data supporting this study’s findings can be found in the GitHub repository (https://github.com/microbemarsh/neuro_gut_axis).

## DECLARATIONS

### Ethics approval and consent to participate

All animal experiments in the study were approved by the Animal Care and Use Committee of Houston Methodist Research Institute, Houston, TX, USA.

### Consent for publication

Not applicable.

### Competing interests

The authors declare that they have no competing interests.

## Notes

### Competing Interest Statement

The authors have declared no competing interest.

## REFERENCES

1. Miller, G.F., et al., Predictors of traumatic brain injury morbidity and mortality: Examination of data from the national trauma data bank: Predictors of TBI morbidity & mortality. Injury, 2021. 52(5): p. 1138–1144.

2. Brett, B.L., et al., Traumatic Brain Injury and Risk of Neurodegenerative Disorder. Biol Psychiatry, 2022. 91(5): p. 498–507.

3. Witcher, K.G., et al., Traumatic Brain Injury Causes Chronic Cortical Inflammation and Neuronal Dysfunction Mediated by Microglia. J Neurosci, 2021. 41(7): p. 1597–1616.

4. Muzio, L., A. Viotti, and G. Martino, Microglia in Neuroinflammation and Neurodegeneration: From Understanding to Therapy. Front Neurosci, 2021. 15: p. 742065.

5. Woodburn, S.C., J.L. Bollinger, and E.S. Wohleb, The semantics of microglia activation: neuroinflammation, homeostasis, and stress. J Neuroinflammation, 2021. 18(1): p. 258.

6. Barbara, G., et al., Inflammatory and Microbiota-Related Regulation of the Intestinal Epithelial Barrier. Front Nutr, 2021. 8: p. 718356.

7. Cryan, J.F., et al., The Microbiota-Gut-Brain Axis. Physiol Rev, 2019. 99(4): p. 1877–2013.

8. Bonaz, B., T. Bazin, and S. Pellissier, The Vagus Nerve at the Interface of the Microbiota-Gut-Brain Axis. Front Neurosci, 2018. 12: p. 49.

9. Sherwin, E., T.G. Dinan, and J.F. Cryan, Recent developments in understanding the role of the gut microbiota in brain health and disease. Ann N Y Acad Sci, 2018. 1420(1): p. 5–25.

10. Chiu, L.S. and R.S. Anderton, The role of the microbiota-gut-brain axis in long-term neurodegenerative processes following traumatic brain injury. Eur J Neurosci, 2023. 57(2): p. 400–418.

11. Parker, B.J., et al., The Genus Alistipes: Gut Bacteria With Emerging Implications to Inflammation, Cancer, and Mental Health. Front Immunol, 2020. 11: p. 906.

12. Hoban, A.E., et al., Regulation of prefrontal cortex myelination by the microbiota. Transl Psychiatry, 2016. 6(4): p. e774.

13. Thion, M.S., et al., Microbiome Influences Prenatal and Adult Microglia in a Sex-Specific Manner. Cell, 2018. 172(3): p. 500–516 e16.

14. Treangen, T.J., et al., Traumatic Brain Injury in Mice Induces Acute Bacterial Dysbiosis Within the Fecal Microbiome. Front Immunol, 2018. 9: p. 2757.

15. Celorrio, M., et al., Gut microbial dysbiosis after traumatic brain injury modulates the immune response and impairs neurogenesis. Acta Neuropathol Commun, 2021. 9(1): p. 40.

16. George, A.K., et al., Rebuilding Microbiome for Mitigating Traumatic Brain Injury: Importance of Restructuring the Gut-Microbiome-Brain Axis. Mol Neurobiol, 2021. 58(8): p. 3614–3627.

17. Huynh, U. and M.L. Zastrow, Metallobiology of Lactobacillaceae in the gut microbiome. J Inorg Biochem, 2023. 238: p. 112023.

18. Opeyemi, O.M., et al., Sustained Dysbiosis and Decreased Fecal Short-Chain Fatty Acids after Traumatic Brain Injury and Impact on Neurologic Outcome. J Neurotrauma, 2021. 38(18): p. 2610–2621.

19. Du, D., et al., Fecal Microbiota Transplantation Is a Promising Method to Restore Gut Microbiota Dysbiosis and Relieve Neurological Deficits after Traumatic Brain Injury. Oxid Med Cell Longev, 2021. 2021: p. 5816837.

20. Lee, J., et al., Gut Microbiota-Derived Short-Chain Fatty Acids Promote Poststroke Recovery in Aged Mice. Circ Res, 2020. 127(4): p. 453–465.

21. Dalile, B., et al., The role of short-chain fatty acids in microbiota-gut-brain communication. Nat Rev Gastroenterol Hepatol, 2019. 16(8): p. 461–478.

22. Martin-Gallausiaux, C., et al., SCFA: mechanisms and functional importance in the gut. Proc Nutr Soc, 2021. 80(1): p. 37–49.

23. Liu, P., et al., The role of short-chain fatty acids in intestinal barrier function, inflammation, oxidative stress, and colonic carcinogenesis. Pharmacol Res, 2021. 165: p. 105420.

24. Sadler, R., et al., Short-Chain Fatty Acids Improve Poststroke Recovery via Immunological Mechanisms. J Neurosci, 2020. 40(5): p. 1162–1173.

25. Tang, C.F., et al., Short-Chain Fatty Acids Ameliorate Depressive-like Behaviors of High Fructose-Fed Mice by Rescuing Hippocampal Neurogenesis Decline and Blood-Brain Barrier Damage. Nutrients, 2022. 14(9).

26. Nagpal, R., et al., Human-origin probiotic cocktail increases short-chain fatty acid production via modulation of mice and human gut microbiome. Sci Rep, 2018. 8(1): p. 12649.

27. Bubnov, R.V., et al., Specific properties of probiotic strains: relevance and benefits for the host. EPMA J, 2018. 9(2): p. 205–223.

28. Hill, C., et al., Expert consensus document. The International Scientific Association for Probiotics and Prebiotics consensus statement on the scope and appropriate use of the term probiotic. Nat Rev Gastroenterol Hepatol, 2014. 11(8): p. 506–14.

29. Villapol, S., D.J. Loane, and M.P. Burns, Sexual dimorphism in the inflammatory response to traumatic brain injury. Glia, 2017. 65(9): p. 1423–1438.

30. Villapol, S., et al., Neurorestoration after traumatic brain injury through angiotensin II receptor blockage. Brain, 2015. 138(Pt 11): p. 3299–315.

31. Villapol, S., K.R. Byrnes, and A.J. Symes, Temporal dynamics of cerebral blood flow, cortical damage, apoptosis, astrocyte-vasculature interaction and astrogliosis in the pericontusional region after traumatic brain injury. Front Neurol, 2014. 5: p. 82.

32. Villapol, S., et al., Smad3 deficiency increases cortical and hippocampal neuronal loss following traumatic brain injury. Exp Neurol, 2013. 250: p. 353–65.

33. Caporaso, J.G., et al., Ultra-high-throughput microbial community analysis on the Illumina HiSeq and MiSeq platforms. ISME J, 2012. 6(8): p. 1621–4.

34. Weisburg, W.G., et al., 16S ribosomal DNA amplification for phylogenetic study. J Bacteriol, 1991. 173(2): p. 697–703.

35. da Veiga Leprevost, F., et al., BioContainers: an open-source and community-driven framework for software standardization. Bioinformatics, 2017. 33(16): p. 2580–2582.

36. Ewels, P.A., et al., The nf-core framework for community-curated bioinformatics pipelines. Nat Biotechnol, 2020. 38(3): p. 276–278.

37. Gruning, B., et al., Bioconda: sustainable and comprehensive software distribution for the life sciences. Nat Methods, 2018. 15(7): p. 475–476.

38. Straub, D., et al., Interpretations of Environmental Microbial Community Studies Are Biased by the Selected 16S rRNA (Gene) Amplicon Sequencing Pipeline. Front Microbiol, 2020. 11: p. 550420.

39. Ewels, P., et al., MultiQC: summarize analysis results for multiple tools and samples in a single report. Bioinformatics, 2016. 32(19): p. 3047–8.

40. Callahan, B.J., et al., DADA2: High-resolution sample inference from Illumina amplicon data. Nat Methods, 2016. 13(7): p. 581–3.

41. Quast, C., et al., The SILVA ribosomal RNA gene database project: improved data processing and web-based tools. Nucleic Acids Res, 2013. 41(Database issue): p. D590–6.

42. Bolyen, E., et al., Reproducible, interactive, scalable and extensible microbiome data science using QIIME 2. Nat Biotechnol, 2019. 37(8): p. 852–857.

43. McMurdie, P.J. and S. Holmes, phyloseq: an R package for reproducible interactive analysis and graphics of microbiome census data. PLoS One, 2013. 8(4): p. e61217.

44. Lin, H. and S.D. Peddada, Multigroup analysis of compositions of microbiomes with covariate adjustments and repeated measures. Nat Methods, 2024. 21(1): p. 83–91.

45. Hamm, R.J., et al., The rotarod test: an evaluation of its effectiveness in assessing motor deficits following traumatic brain injury. J Neurotrauma, 1994. 11(2): p. 187–96.

46. Villapol, S., et al., Candesartan, an angiotensin II AT(1)-receptor blocker and PPAR-gamma agonist, reduces lesion volume and improves motor and memory function after traumatic brain injury in mice. Neuropsychopharmacology, 2012. 37(13): p. 2817–29.

47. Pellow, S. and S.E. File, Anxiolytic and anxiogenic drug effects on exploratory activity in an elevated plus-maze: a novel test of anxiety in the rat. Pharmacol Biochem Behav, 1986. 24(3): p. 525–9.

48. Porsolt, R.D., et al., Behavioural despair in rats: a new model sensitive to antidepressant treatments. Eur J Pharmacol, 1978. 47(4): p. 379–91.

49. Hemarajata, P. and J. Versalovic, Effects of probiotics on gut microbiota: mechanisms of intestinal immunomodulation and neuromodulation. Therap Adv Gastroenterol, 2013. 6(1): p. 39–51.

50. Martin, B.D., D. Witten, and A.D. Willis, Modeling Microbial Abundances and Dysbiosis with Beta-Binomial Regression. Ann Appl Stat, 2020. 14(1): p. 94–115.

51. Cryan, J.F., et al., The gut microbiome in neurological disorders. Lancet Neurol, 2020. 19(2): p. 179–194.

52. Soriano, S., et al., Fecal Microbiota Transplantation Derived from Alzheimer’s Disease Mice Worsens Brain Trauma Outcomes in Wild-Type Controls. Int J Mol Sci, 2022. 23(9).

53. Ullah, H., et al., The gut microbiota-brain axis in neurological disorder. Front Neurosci, 2023. 17: p. 1225875.

54. Cannon, A.R., et al., Traumatic Brain Injury-Induced Inflammation and Gastrointestinal Motility Dysfunction. Shock, 2023. 59(4): p. 621–626.

55. Zheng, Z., et al., Gut Microbiota Dysbiosis after Traumatic Brain Injury Contributes to Persistent Microglial Activation Associated with Upregulated Lyz2 and Shifted Tryptophan Metabolic Phenotype. Nutrients, 2022. 14(17).

56. Urban, R.J., et al., Altered Fecal Microbiome Years after Traumatic Brain Injury. J Neurotrauma, 2020. 37(8): p. 1037–1051.

57. Crovesy, L., D. Masterson, and E.L. Rosado, Profile of the gut microbiota of adults with obesity: a systematic review. Eur J Clin Nutr, 2020. 74(9): p. 1251–1262.

58. Ma, Y., et al., Lactobacillus acidophilus Exerts Neuroprotective Effects in Mice with Traumatic Brain Injury. J Nutr, 2019. 149(9): p. 1543–1552.

59. Wan, G., et al., Effects of probiotics combined with early enteral nutrition on endothelin-1 and C-reactive protein levels and prognosis in patients with severe traumatic brain injury. J Int Med Res, 2020. 48(3): p. 300060519888112.

60. Gamallat, Y., et al., Lactobacillus rhamnosus induced epithelial cell apoptosis, ameliorates inflammation and prevents colon cancer development in an animal model. Biomed Pharmacother, 2016. 83: p. 536–541.

61. Alipour Nosrani, E., et al., Neuroprotective effects of probiotics bacteria on animal model of Parkinson’s disease induced by 6-hydroxydopamine: A behavioral, biochemical, and histological study. J Immunoassay Immunochem, 2021. 42(2): p. 106–120.

62. Authier, H., et al., Oral Administration of Lactobacillus helveticus LA401 and Lactobacillus gasseri LA806 Combination Attenuates Oesophageal and Gastrointestinal Candidiasis and Consequent Gut Inflammation in Mice. J Fungi (Basel), 2021. 7(1).

63. Eslami, M., et al., Probiotics function and modulation of the immune system in allergic diseases. Allergol Immunopathol (Madr), 2020. 48(6): p. 771–788.

64. Vemuri, R., et al., Lactobacillus acidophilus DDS-1 Modulates Intestinal-Specific Microbiota, Short-Chain Fatty Acid and Immunological Profiles in Aging Mice. Nutrients, 2019. 11(6).

65. Zaydi, A.I., et al., Lactobacillus plantarum DR7 improved brain health in aging rats via the serotonin, inflammatory and apoptosis pathways. Benef Microbes, 2020. 11(8): p. 753–766.

66. Wang, W., et al., Lactobacillus plantarum Combined with Galactooligosaccharides Supplement: A Neuroprotective Regimen Against Neurodegeneration and Memory Impairment by Regulating Short-Chain Fatty Acids and the c-Jun N-Terminal Kinase Signaling Pathway in Mice. J Agric Food Chem, 2022. 70(28): p. 8619–8630.

67. Chong, H.X., et al., Lactobacillus plantarum DR7 alleviates stress and anxiety in adults: a randomised, double-blind, placebo-controlled study. Benef Microbes, 2019. 10(4): p. 355–373.

68. Slykerman, R.F., et al., Effect of Lactobacillus rhamnosus HN001 in Pregnancy on Postpartum Symptoms of Depression and Anxiety: A Randomised Double-blind Placebo-controlled Trial. EBioMedicine, 2017. 24: p. 159–165.

69. Sun, B., et al., The Effects of Lactobacillus acidophilus on the Intestinal Smooth Muscle Contraction through PKC/MLCK/MLC Signaling Pathway in TBI Mouse Model. PLoS One, 2015. 10(6): p. e0128214.

70. Wang, J.J., et al., Modulatory effect of Lactobacillus acidophilus KLDS 1.0738 on intestinal short-chain fatty acids metabolism and GPR41/43 expression in beta-lactoglobulin-sensitized mice. Microbiol Immunol, 2019. 63(8): p. 303–315.

71. Grazul, H., L.L. Kanda, and D. Gondek, Impact of probiotic supplements on microbiome diversity following antibiotic treatment of mice. Gut Microbes, 2016. 7(2): p. 101–14.

72. Xiao, W., et al., Correction to: The microbiota-gut-brain axis participates in chronic cerebral hypoperfusion by disrupting the metabolism of short-chain fatty acids. Microbiome, 2022. 10(1): p. 70.

73. Church, J.S., et al., Serum short chain fatty acids mediate hippocampal BDNF and correlate with decreasing neuroinflammation following high pectin fiber diet in mice. Front Neurosci, 2023. 17: p. 1134080.

74. Caetano-Silva, M.E., et al., Inhibition of inflammatory microglia by dietary fiber and short-chain fatty acids. Sci Rep, 2023. 13(1): p. 2819.

75. Erny, D., et al., Host microbiota constantly control maturation and function of microglia in the CNS. Nat Neurosci, 2015. 18(7): p. 965–77.

76. He, J., et al., Short-Chain Fatty Acids and Their Association with Signalling Pathways in Inflammation, Glucose and Lipid Metabolism. Int J Mol Sci, 2020. 21(17).

77. Portincasa, P., et al., Gut Microbiota and Short Chain Fatty Acids: Implications in Glucose Homeostasis. Int J Mol Sci, 2022. 23(3).

78. Spichak, S., et al., Microbially-derived short-chain fatty acids impact astrocyte gene expression in a sex-specific manner. Brain Behav Immun Health, 2021. 16: p. 100318.

79. Shastri, P., et al., Sex differences in gut fermentation and immune parameters in rats fed an oligofructose-supplemented diet. Biol Sex Differ, 2015. 6: p. 13.

80. Sun, Z., et al., Living, Heat-Killed Limosilactobacillus mucosae and Its Cell-Free Supernatant Differentially Regulate Colonic Serotonin Receptors and Immune Response in Experimental Colitis. Nutrients, 2024. 16(4).

81. Guo, P., et al., Clostridium species as probiotics: potentials and challenges. J Anim Sci Biotechnol, 2020. 11: p. 24.

82. Wang, S., et al., Ingestion of Faecalibaculum rodentium causes depression-like phenotypes in resilient Ephx2 knock-out mice: A role of brain-gut-microbiota axis via the subdiaphragmatic vagus nerve. J Affect Disord, 2021. 292: p. 565–573.

83. Hassan, M.U., et al., Probiotic Properties of Lactobacillus helveticus and Lactobacillus plantarum Isolated from Traditional Pakistani Yoghurt. Biomed Res Int, 2020. 2020: p. 8889198.

84. Edgar, R.C., Accuracy of taxonomy prediction for 16S rRNA and fungal ITS sequences. PeerJ, 2018. 6: p. e4652.

85. Durazzi, F., et al., Comparison between 16S rRNA and shotgun sequencing data for the taxonomic characterization of the gut microbiota. Sci Rep, 2021. 11(1): p. 3030.

86. Johnson, J.S., et al., Evaluation of 16S rRNA gene sequencing for species and strain-level microbiome analysis. Nat Commun, 2019. 10(1): p. 5029.

87. Tan, M., et al., Effects of probiotics on serum levels of Th1/Th2 cytokine and clinical outcomes in severe traumatic brain-injured patients: a prospective randomized pilot study. Crit Care, 2011. 15(6): p. R290.

88. Sun, J., et al., Clostridium butyricum attenuates cerebral ischemia/reperfusion injury in diabetic mice via modulation of gut microbiota. Brain Res, 2016. 1642: p. 180–188.

89. Yang, X., et al., Gut microbiota mediates the protective role of Lactobacillus plantarum in ameliorating deoxynivalenol-induced apoptosis and intestinal inflammation of broiler chickens. Poult Sci, 2020. 99(5): p. 2395–2406.

90. Davoli-Ferreira, M., C.A. Thomson, and K.D. McCoy, Microbiota and Microglia Interactions in ASD. Front Immunol, 2021. 12: p. 676255.

91. Jiang, C.T., et al., Modulators of microglia activation and polarization in ischemic stroke (Review). Mol Med Rep, 2020. 21(5): p. 2006–2018.

92. Baudo, G., et al., Sex-dependent improvement in traumatic brain injury outcomes after liposomal delivery of dexamethasone in mice. bioRxiv, 2023.

93. Zinger, A., et al., Biomimetic Nanoparticles as a Theranostic Tool for Traumatic Brain Injury. Adv Funct Mater, 2021. 31(30): p. 2100722.

94. Gareau, M.G., et al., Probiotics prevent death caused by Citrobacter rodentium infection in neonatal mice. J Infect Dis, 2010. 201(1): p. 81–91.

95. Beilharz, J.E., et al., Cafeteria diet and probiotic therapy: cross talk among memory, neuroplasticity, serotonin receptors and gut microbiota in the rat. Mol Psychiatry, 2018. 23(2): p. 351–361.

96. Steenbergen, L., et al., A randomized controlled trial to test the effect of multispecies probiotics on cognitive reactivity to sad mood. Brain Behav Immun, 2015. 48: p. 258–64.

