## Supplementary figures and images for "Probiotic treatment causes sex-specific neuroprotection after traumatic brain injury in mice"

### Supplementary Figure 1

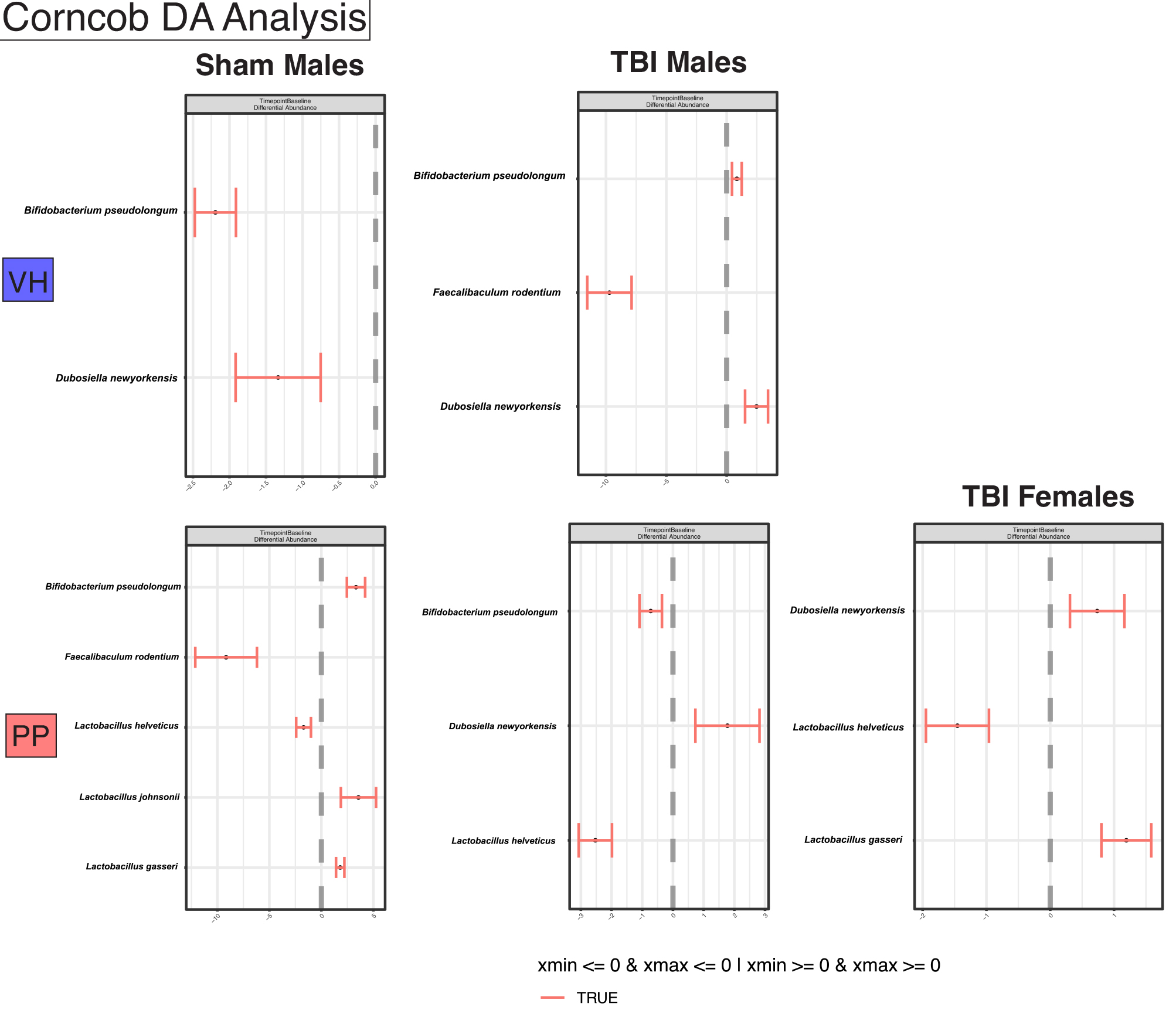
